# An Inducible CRISPRi system for phenotypic analysis of essential genes in *Pseudomonas aeruginosa*

**DOI:** 10.1101/2025.09.11.675340

**Authors:** Jaryd R. Sullivan, Kristina M. Ferrara, Rebecca Barrick, Keith P. Romano, Thulasi Warrier, Deborah T. Hung

## Abstract

Precise and tunable genetic tools are essential for high throughput functional genomics. To address this need in the important gram-negative pathogen *Pseudomonas aeruginosa*, we developed and characterized a tightly regulated CRISPRi system that enables precise and tunable repression of essential genes. The system utilizes a rhamnose-inducible promoter to control both the *Streptococcus pasteurianus*-derived dCas9 and gene-specific sgRNAs, each encoded on separate plasmids for modularity and efficiency. The combination of tight regulation and high conjugation efficiency facilitated the rapid and facile construction of strains with regulated depletion of 16 essential genes spanning diverse pathways. Comparison of phenotypes across the different genetically depleted strains, including growth rate, susceptibility to antibiotics, and changes in transcriptional programs, revealed novel aspects of gene function or small molecule mechanism of action. Finally, the rhamnose-inducible CRISPRi system supports the generation and stable maintenance of pooled mutant libraries, thereby paving the way for future genome-wide, systematic assessment of individual gene vulnerabilities, which will provide critical insights for target prioritization in antibiotic discovery efforts against this recalcitrant pathogen.

**IMPORTANCE:** CRISPRi has become an invaluable tool for studying genetics. In particular, the ability to knockdown genes enables the study of essential genes and their role in cell survival. However, a tightly regulated gene knockdown system is required to gain valuable insights into these vulnerable genes by virtue of their essentiality. We report a tightly regulated CRISPRi system to study the biology of essential gene perturbations in *Pseudomonas aeruginosa*, an important gram-negative pathogen that causes severe infections and is increasingly resistant to current antibiotics. This system enables characterization of both chemical genetic interactions between small molecules and specific gene depletions, and the impact of genetic perturbations on transcriptional networks. Genetic perturbations using CRISPRi can thus further our understanding of basic biology with translation towards future antimicrobial development.

## INTRODUCTION

Bacterial genetics has a long history of enabling the elucidation of individual gene roles in critical cellular functions and complex phenotypes that shape microbial physiology, pathogenesis, and antibiotic resistance. In the past, genetic strategies for gene disruption have included targeted allelic exchange with two-step homologous recombination, phage recombinase-based single step replacement with marker genes, as well as non-targeted approaches using transposon insertion mutagenesis for gene disruption.^1–3^ Although well- suited for studying non-essential genes that are dispensable for bacterial growth, these approaches are more cumbersome when applied to essential genes where graded depletion rather than total disruption is often desired. Genetic strategies for essential gene depletion have historically required replacing the native promoter with an inducible promoter or inserting protein degradation tags to control the level of expression.^4–6^ Recently however, the advent of CRISPR-interference (CRISPRi) technology has transformed the ability to perturb essential genes in a scalable fashion.^7,8^ By programming a catalytically inactive Cas9 (dCas9) to bind—but not cut—specific DNA sequences guided by an sgRNA, transcription of the target gene can be silenced through steric hindrance that blocks RNA polymerase at the site.

CRISPRi has proven to be a valuable resource in the field of bacterial genetics with well-characterized systems developed for model organisms like *Escherichia coli*^7,9,10^, *Bacillus subtilis*^11^, and *Streptococcus pneumoniae*^12^ with the power to study the biological functions of essential genes and perform CRISPRi screens. A mobile CRISPRi system was created to further facilitate rapid CRISPRi implementation into a variety of other gram-positive and negative bacteria with an inducible *dcas9* from *Streptococcus pyogenes* and constitutively expressed sgRNA.^13^ While this *S. pyogenes* dCas9 based system was less effective in *P. aeruginosa* due to observed instability of the dCas9 protein^13^, a more optimized *S. pyogenes* system was recently reported^14^. A different CRISPRi tool based on *Streptococcus pasteurianus* dCas9 has also been reported in *Pseudomonas spp*. with up to 1000-fold reduction in targeted activity, but the expression of *dcas9* from anhydrotetracycline or IPTG inducible promoters was leaky, which makes targeting essential genes more challenging.^15^ Activation of the system in the absence of inducer can preclude the construction, isolation, and thus study of depletion mutants corresponding to essential genes due to uncontrolled target depletion leading to a constitutive fitness cost. This fitness cost can result in selection for suppressor mutations just from simple propagation of the hypomorphic strain, can preclude the shotgun generation of comprehensive, pooled, genome-wide libraries targeting essential genes as less fit mutants will be competed out by more fit ones, or can preclude construction of the strain altogether. In *Mycobacteria spp*., this problem was solved by tightly controlling both *dcas9* from *Streptococcus thermophilus* and sgRNA expression from the engineered anhydrotetracycline-driven promoters.^16^ This advancement enabled the successful evaluation not simply of gene essentiality but also the relative vulnerability of the essential genes on a genome-wide scale.^17^ A similar strategy with an engineered tetracycline-inducible promoter aided optimization of the *S. pyogenes* dCas9-based tool in *P. aeruginosa*.^14^

Here, we developed a tightly regulated CRISPRi system with the *S. pasteurianus* dCas9 to enable phenotypic profiling of individual essential genes or pooled CRISPRi studies in *P. aeruginosa*, a WHO- prioritized pathogen for urgent antimicrobial discovery.^18^ It is based on a two-piece modular strategy with dual inducible-promoter control of the catalytically inactive dCas9 and the sgRNA. Unlike many of the CRISPRi systems that rely on induction of *dcas9* and constitutive expression of the sgRNA, we utilized the rhamnose-inducible P_rhaBAD_ promoter for both *dcas9* and sgRNA expression.^19–21^ This system provided tight control with increased knockdown efficiency in the presence of inducer and minimal knockdown in the absence of inducer, thus facilitating detailed functional characterization of essential genes. In addition, we separated *dcas9* from the sgRNA which enabled the generation of a parental *P. aeruginosa*::*dcas9* strain into which smaller sgRNA-carrying plasmids could be integrated with greater success, thereby enabling construction of pooled CRISPRi libraries targeting essential genes rapidly and with ease. Finally, we demonstrated the tunability of this tool both by partially depleting antibiotic targets to explore chemical genetic interactions with anti-pseudomonal drugs, and by maximally depleting essential genes to identify perturbation-specific transcriptional responses in *P. aeruginosa*.

## RESULTS

### An inducible CRISPRi system in *Pseudomonas aeruginosa* using *S. pasteurianus* dCas9

To develop a tightly regulated CRISPRi system for essential gene perturbations in *P. aeruginosa*, we chose to focus on systems with inducers that are minimally impacted by efflux pumps in *P. aeruginosa* to preserve their utility in a wide breadth of strains including clinical isolates that often display hyper-efflux phenotypes.^22,23^ We started with the arabinose inducible P_araBAD_ promoter to drive *dcas9*, since the IPTG-inducible P_lac_/P_tac_ promoters had been reported to cause leaky CRISPRi activation in *P. aeruginosa*.^15^ We paired arabinose-inducible *dcas9* expression with a constitutive P_ProD_ promoter driving sgRNA expression (termed CRISPRi^A^, for arabinose). Both were incorporated into a single *att*Tn7 site-integrating mini-Tn7 vector^24^ and introduced into *P. aeruginosa* strain PA14 (Figure 1A). When we compared codon optimized, His-tagged *dcas9* variants^15,16,25,26^, including dCas9 from *S. pasteurianus* (dCas9_Spa_), *S. pyogenes* (dCas9_Spy_), and *S. thermophilus* (dCas9_Sth1_), we found by Western blot analysis using an anti-His antibody that dCas9_Spa_ showed the strongest arabinose dose-dependent expression levels, followed by dCas9_Spy_, while there were no detectable levels of dCas9_Sth1_ (Figure 1B).

**Figure 1.**
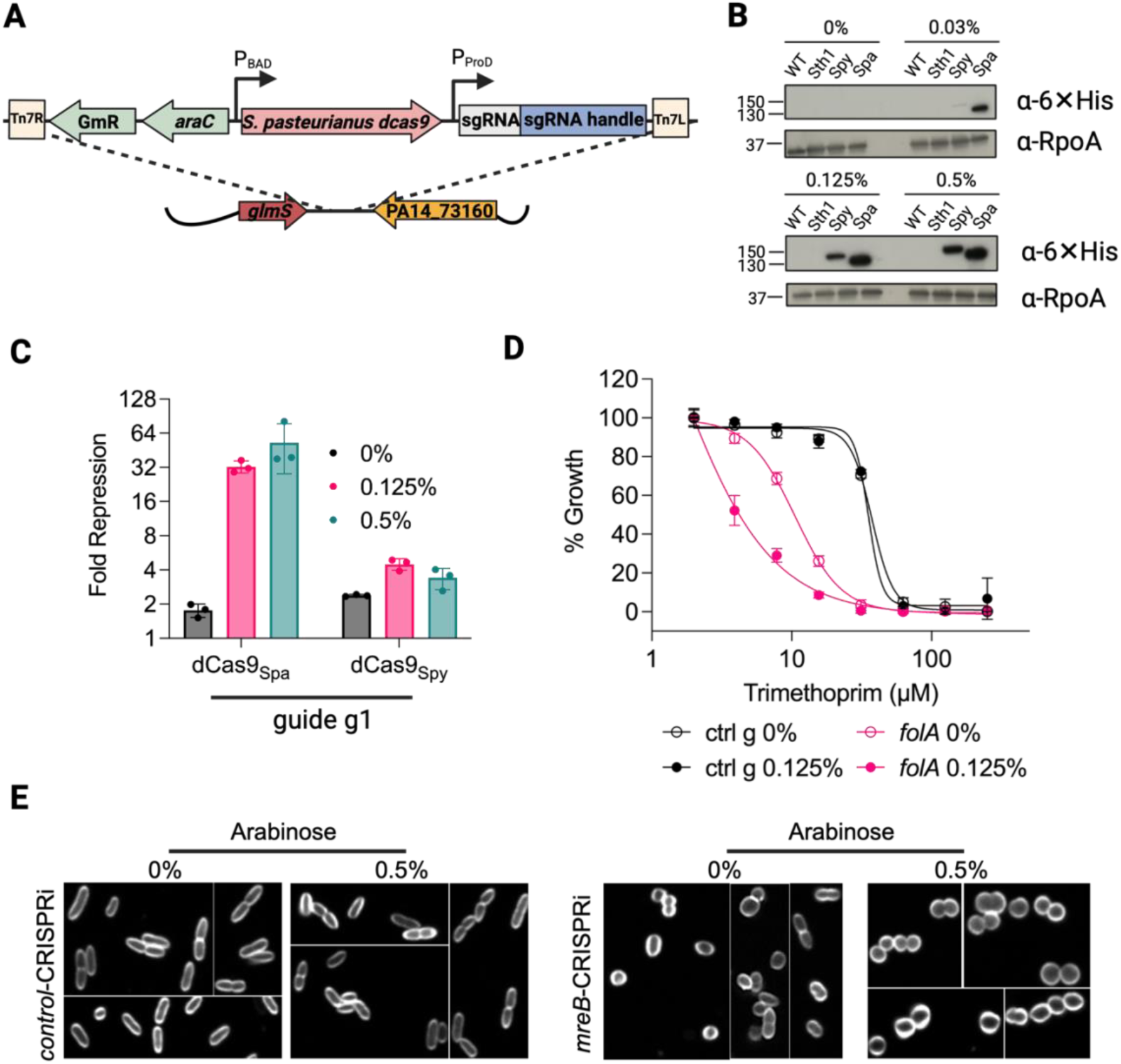
An arabinose-based CRISPRi system yields leaky CRISPRi phenotypes. (A) Schematic of CRISPRi P_araBAD_ system (CRISPRi^A^). dCas9 expression is driven by the arabinose-inducible promoter P_araBAD_ with sgRNA constitutively expressed under the P_ProD_ promoter. CRISPRi^A^ system integrates at the *att*Tn7 site in the PA14 genome. (B) Western Blot analysis of expression of C-terminal 6XHis-tagged dCas9 variants in PA14 strain at varying arabinose concentrations. Wild-type PA14 or PA14 expressing dCas9 from *S. thermophilus* (Sth1), *S. pyogenes* (Spy), or *S. pasteurianus* (Spa) was lysed after induction with 0, 0.03, 0.125, or 0.5% arabinose and blotted with an anti-His antibody. dCas9_Spa_ and dCas9_Spy_ levels were arabinose dose-dependent with higher abundance of dCas9_Spa_ while dCas9_Sth1_ was not expressed. (C) mKate2 fluorescence repression (normalized to control sgRNA) by CRISPRi^A^ using dCas9_Spa_ or dCas9_Spy_ with sgRNA g1 (Figures S1A) measured after 15 hours of growth in M9 minimal media with 0, 0.125, or 0.5% arabinose. dCas9_Spa_ induction led to stronger depletion of mKate2 than dCas9_Spy_, whose activity was also not arabinose dependent. (D) Sensitization of *folA*-CRISPRi^A^ to trimethoprim. Activity of trimethoprim against CRISPRi^A^ strains with control or *folA1* (Figure S1H) sgRNAs grown in LB broth with and without 0.125% arabinose for 24 hours. Trimethoprim sensitization was dependent on having the *folA1* guide but independent of arabinose. (E) Morphology of CRISPRi^A^ strains with control or *mreB* sgRNAs. CRISPRi^A^ strains were grown in 0% or 0.5% arabinose for ∼19h, stained with FM-143x, and visualized using confocal microscopy. *mreB-*CRISPRi^A^ cells displayed spherical morphology even in the absence of arabinose.

To compare gene knockdown efficiency, we generated a reporter system based on the fluorescent protein mKate2^27,28^ and designed guides to the promoter and along the length of the *mkate2* coding sequence (Figure S1A). Not only did dCas9_Spa_ repress mKate2 fluorescence more than dCas9_Spy_ (∼50- vs 4- fold), but it showed both tighter control in the absence of inducer and greater inducer dose-dependence than the *S. pyogenes* based system (Figure 1C and S1B). Western blot analysis revealed that dCas9_Spy_ likely undergoes some degree of degradation in *P. aeruginosa* as evidenced at higher arabinose concentrations (Figure S1C).^13^ We therefore focused on the dCas9_Spa_-based system for further characterization and optimization, consistent with a previous report using dCas9_Spa_.^15^

### Expanding the functional PAM variants for dCas9_Spa_ in *P. aeruginosa*

Many factors, including proto-adjacent spacer motif (PAM) sequence, guide sequence location within the gene, guide length, and template versus non-template strand targeting, influence CRISPRi efficiency.^7^ Three dCas9_Spa_-specific PAM sequences have been previously reported.^15^ To try to expand the number of PAM sites that could be targeted in the *P. aeruginosa* genome, we systematically explored variations in PAM sequence using the mKate2 reporter assay. We tested the three previously reported dCas9_Spa_-specific PAM sequences^15^ (NNGTGA, NNGCGA, NNGTAA) and seven additional variants to expand the range of the guide sequence and location. Previous sequences NNGTGA and NNGCGA indeed provided the highest degrees of fluorescence repression (∼8 to 64-fold) irrespective of guide location (Figure S1D). On the other hand, while NNGTAA generally exhibited lower repression (∼2 to 8-fold), its activity was location dependent with the guide located 48bp into the open reading frame of mKate2 (g5) providing the strongest repression of all tested guides (Figure S1D). We identified a new PAM, NNGTAT, with efficiency identical to the known NNGTAA, thus expanding the repertoire of dCas9_Spa_-specific PAMs and the number of PAM sites in the *P. aeruginosa* genome by an additional 10% (Figure S1E). As expected, changing the first specific nucleotide or more than one nucleotide in these PAM sequences, when introduced in the *mKate2* transcription start site, completely abrogated dCas9 activity; similarly targeting the template strand resulted in no activity, as evidenced by the lack of activity of a guide designed to do so (sgRNA g12; Figure S1D).^16,29^ Notably, variability in the length of the specificity region of the sgRNA from the standard 21-24nt to 16nt did not alter knockdown efficiency (Figure S1F). Based on these findings, we prioritized the three known PAMs during guide design to select 20-24nt long guides to target the non-template strand of genes of interest at sites most proximal to the transcriptional start site. Although we defined a new PAM, NNGTAT, it was not included due to its reduced strength.

### Activity of the arabinose-dependent dCas9_Spa_ system occurred in the absence of inducer

Next, we evaluated the system’s efficacy towards endogenous essential genes in PA14 strain. Targeting the dihydrofolate reductase gene, *folA*, with three sgRNAs that bound different locations led to variable growth defects compared to the *control*-CRISPRi^A^ strain (Figure S1G). While in the absence of arabinose, all three strains grew normally, in the presence of arabinose, guide 1 located in the promoter region led to the strongest growth deficit, guide 2 located 28 bp into the open reading frame had a milder impact, and guide 3 located 159 bp into the open reading frame had no effect (Figure S1H). However, while normal strain growth in the absence of inducer was encouraging, in a different, more sensitive assay in which we measured the MIC_90_ of a FolA inhibitor (trimethoprim) in the CRISPRi^A^ mutants, we observed lower MIC_90_ values in the *folA*- CRISPRi^A^ strain relative to the *control*-CRISPRi^A^ strain, even in the absence of inducer (Figure 1D). This increased trimethoprim sensitivity implied that there was some baseline level of FolA depletion despite the apparent normal growth, indicating leaky activation of the CRISPRi^A^ system. The observed leakiness was not just isolated to *folA*. We similarly observed phenotypic consequences in the absence of inducer in a strain designed to deplete *mreB*, which controls the rod shape of gram-negative bacteria. In uninduced conditions, *mreB*-CRISPRi^A^ bacteria also had normal growth based on optical density (OD_600_); however, they had a spherical morphology, in contrast to the typical rod shape of wild-type bacteria and the *control*-CRISPRi^A^ strain, suggesting some degree of target depletion (Figure 1E).

### Development of a stronger and tighter CRISPRi system in *P. aeruginosa*

To create a more tightly regulated system, we substituted the arabinose P_araBAD_ promoter with the rhamnose-inducible P_rhaBAD_ promoter, which is known to be more tightly regulated in *P. aeruginosa*.^19,30^ Additionally, we used the P_rhaBAD_ promoter to regulate expression of both *dcas9* and the guide. We designed it as a two-plasmid system to make it more modular and avoid cloning inefficiency due to the large size of the mini-Tn7 vector when the P_rhaBAD_ promoter elements were introduced. The *dcas9*_Spa_ was integrated at the *ϕCTX* attachment site in wild-type PA14 genome, thereby generating a parental CRISPRi strain, (PA14::*dcas9)*. Different sgRNAs were then separately and easily integrated at the *att*Tn7 site using a smaller mini-Tn7 plasmid (Figure 2A). Knockdown efficiency of this P_rhaBAD_-based system (termed CRISPRi^R^, for rhamnose) was comparable to the P_araBAD_ system on mKate2 repression (Figure 2B); however, activation of the system was now tightly dependent on rhamnose, as evidenced by the strict rhamnose-dependence of trimethoprim sensitization of *folA*-CRISPRi^R^ and the absence of morphological changes observed in *mreB*-CRISPRi^R^ strains in the absence of rhamnose (Figures 2C and 2D).

**Figure 2.**
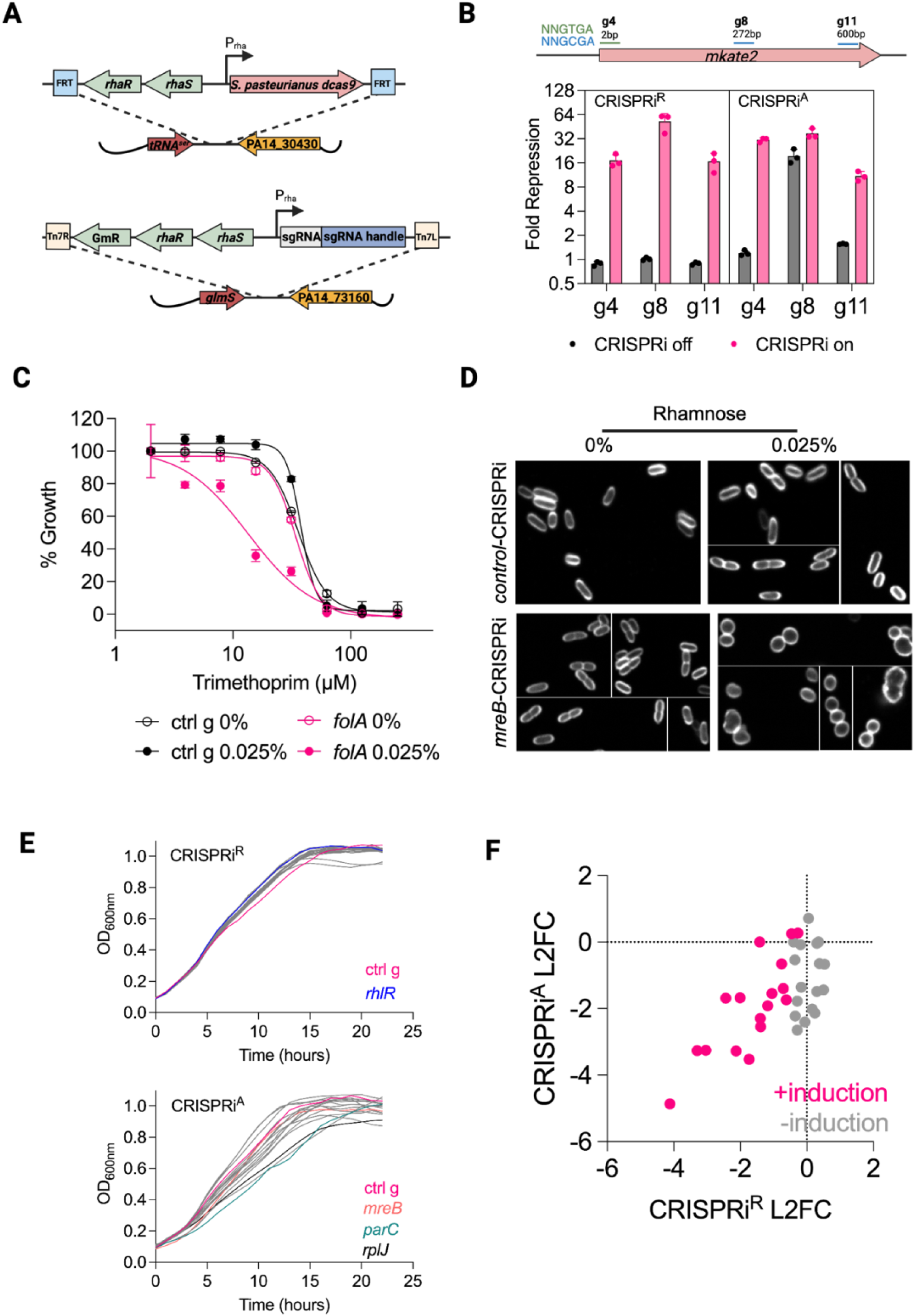
A rhamnose-based, modular two-piece CRISPRi P_rhaBAD_ system (CRISPRi^R^) eliminates leaky phenotypes. (A) Schematic of CRISPRi^R^ system. dCas9_Spa_ expression is driven by the rhamnose-inducible promoter P_rhaBAD_ and is integrated at the *Φ*CTX site in the PA14 genome. This establishes a parental dCas9_Spa_ expressing strain of PA14. sgRNAs are also under control of the P_rhaBAD_ promoter and is integrated at the *att*Tn7 site in the PA14 genome. (B) In the presence of inducer, CRISPRi^R^ activity against the mKate2 reporter is comparable to CRISPRi^A^, but in the absence of inducer, CRISPRi^R^ is more tightly off. mKate2 fluorescence repression (normalized to control sgRNA) by CRISPRi with sgRNAs g4, g8, and g11 measured after 15 hours of growth in minimal M9 media with 0.05% rhamnose (CRISPRi^R^) or 0.5% arabinose (CRISPRi^A^). (C) Sensitization of *folA*-CRISPRi^R^ to trimethoprim. Activity of trimethoprim against CRISPRi^R^ strains with control or *folA1* sgRNAs (Figure S1H) grown in LB broth for 24 hours. Trimethoprim sensitization was dependent on expressing the *folA1* sgRNA and on the presence of rhamnose. (D) Morphology of CRISPRi^R^ strains with control or *mreB* sgRNAs. CRISPRi^R^ strains were grown in 0% or 0.05% rhamnose for ∼19h, stained with FM-143x, and visualized using confocal microscopy. *mreB*-CRISPRi^R^ cells displayed spherical morphology only in the presence of the CRISPRi^R^ inducer indicating a tight off state. (E) Comparison of CRISPRi growth curves in LB broth without inducer. CRISPRi^R^ strains targeting essential genes exhibited uniform growth in the absence of rhamnose while CRISPRi^A^ strains exhibited variable growth in the absence of arabinose. Each line is the growth curve for a different CRISPRi strain with the control-CRISPRi strain shown in pink. (F) Increased control of the CRISPRi^R^ system limits leaky knockdown of target genes. CRISPRi knockdown of target genes analyzed by RNA-seq of each corresponding CRISPRi depletion strain. The degree of knockdown of the targeted gene is quantified using log_2_ fold change (L2FC) relative to *control*-CRISPRi strain after 6 hours of growth in 0.5% arabinose (CRISPRi^A^) or 0.05% rhamnose (CRISPRi^R^). Each dot represents one CRISPRi strain either in induced (pink) or uninduced conditions (grey). Without induction, there is no decrease in expression of the CRISPRi^R^ targeted gene in the corresponding strain, in contrast to the CRISPRi^A^ system.

### Targeting an expanded set of endogenous genes in *P. aeruginosa*

We compared the new rhamnose-based system with the arabinose system by individually constructing, in parallel, 16 genes that are part of the essential core genome of *P. aeruginosa*.^31^ Selected genes are involved in DNA synthesis (*folA*, *gyrA*, *gyrB*, *parC*), protein synthesis (*rpoB*, *rpsL*, *rplJ*, *leuS*), and cell envelope biogenesis (*lptA*, *lptD*, *lptE*, *dxr*, *lpxC*, *mreB*, *murA*, *pbpA*). We included the non-essential transcription factor *rhlR*, which coordinates pyocyanin production as a negative control with knockdown resulting in visible reduction in cyan-pigmented pyocyanin in the medium in a rhamnose dose dependent manner (Figure S2A). The tightness of the new CRISPRi^R^ system was validated by observing more uniform growth curves across the 18 CRISPRi strains in the ‘off’ state compared to the leakier CRISPRi^A^ system (Figure 2E). Enumeration of transcript levels of each target gene by RNA-seq showed that there was no gene depletion in the corresponding CRISPRi strain using the CRISPRi^R^ system in the absence of rhamnose in contrast to measurable gene depletion in the corresponding strain using the CRISPRi^A^ system in the absence of arabinose (Figure 2F). We also compared the regulation of the arabinose and rhamnose CRISPRi systems by propagating the 18 CRISPRi strains uninduced in a pooled fashion. CRISPRi strains were grown separately without inducer and pooled at equal density and continuously passaged without inducer for up to 10h (Figure S3A). We sampled the pools over time, amplified the sgRNAs from the pool, and sequenced the amplicon pools to enumerate the surviving census of each strain in the pool. We observed dropout of certain strains over time using the CRISPRi^A^ system, which confirmed reduced fitness of some strains due to leaky downregulation of their corresponding targets even in the absence of inducer (Figure 3A). On the other hand, pool diversity with equal representation of each strain was maintained with the CRISPRi^R^ system (Figure 3A).

**Figure 3.**
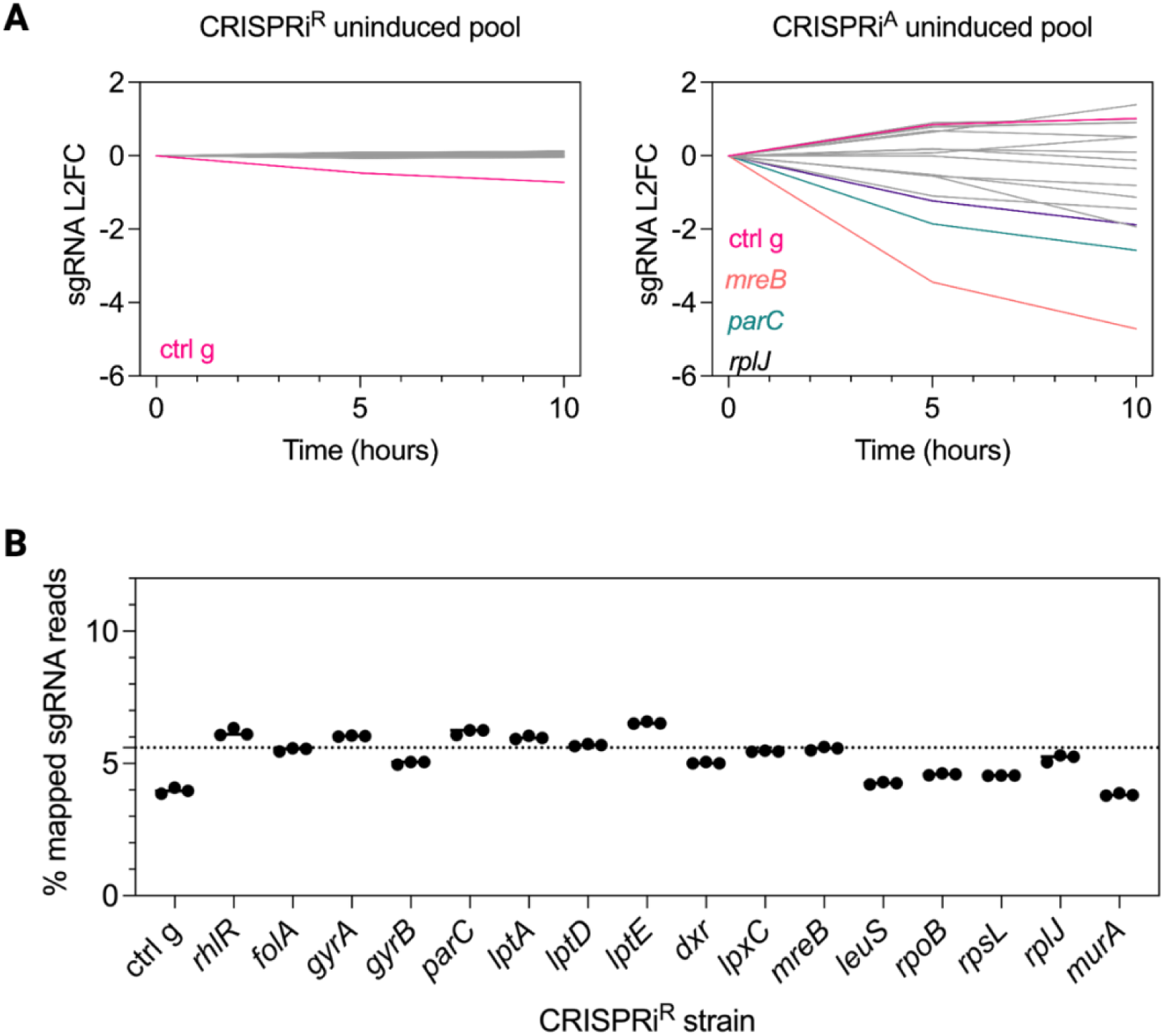
Tight control of CRISPRi^R^ facilitates pooled CRISPRi library construction and propagation. (A) Targeted amplicon sequencing of sgRNAs as a proxy for individual strain census within a propagated pool of CRISPRi^R^ or CRISPRi^A^ strains without inducer. 18 individual CRISPRi strains were pooled at equal ratios at time zero. No loss of strains was detected in the CRISPRi^R^ pool even after 10h of growth, as determined by log_2_ fold-change (L2FC) in the sgRNA amplicon abundance corresponding to the engineered strain targeting each essential gene (left). In contrast, strain dropout was apparent in the pool of CRISPRi^A^ strains (right). Each line represents a different CRISPR mutant, with colored lines referring to the corresponding colored gene shown. (B) Pooled construction of a single CRISPRi^R^ library containing 17 different strains. Conjugation was carried out with a pool of 17 different *E. coli* sgRNA donor strains (pooled at equal ratios). Targeted amplicon sequencing to measure the abundance of sgRNA as a proxy for strain abundance is shown as a percentage of all mapped sgRNA reads (dotted line indicates 5.8% theoretical target for each CRISPRi^R^ sgRNA). All the individual strains were recovered at the expected ∼5% of the total pool abundance.

To further support the benefit of a tightly regulated system, we were able to not simply propagate but also directly generate in a single pool, a library of CRISPRi^R^ strains targeting these essential genes (Figure S3B). First, we cloned sgRNAs individually into the miniTn7 sgRNA vector in the *E. coli* donor strain. Second, we set up a tetrapartite mating strategy with recipient PA14::*dcas9* strain, two *E. coli* helper strains harboring the helper plasmids pRK2013 and pTNS2, respectively, and a pool of *E. coli* donor strains mixed at equal proportions carrying the CRISPRi^R^ sgRNA vectors. After selecting for successful PA14 conjugants on irgasan and gentamicin, all colonies were pooled together for targeted amplicon sequencing of the guides. With 17 different sgRNAs in the pool, we expected each sgRNA to make up 5.8% of all total mapped reads (dashed grey line) and indeed, we obtained ∼5% mapped sgRNA reads for each sgRNA (Figure 3B). The tightly regulated rhamnose system thus not only minimized loss of CRISPRi^R^ strain diversity with propagation over time but enabled construction of a strain pool with equal representation of individual CRISPRi^R^ strains.

### Growth phenotypes of essential genes depleted in *P. aeruginosa*

Upon activation of the CRISPRi^R^ machinery, we observed a range of gene knockdown levels and growth phenotypes across the different gene targets, despite selecting optimal PAMs which were most proximal to the transcription/translation start site and using one of the three strongest PAMs for all genes (Figure S2B). The relationship between gene identity, degree of knockdown, and impact on phenotypes including growth is complicated, captured in the concept of gene vulnerability and the sensitivity of the cell to target depletion.^17,32^ We observed variable growth phenotypes even when knocking down targets involved in the same pathway or function.

We observed that gene depletion can lead to distinct phenotypic outcomes, ranging from bactericidal effects to bacteriostatic behavior. When we knocked down *murA*, which is involved in peptidoglycan synthesis, we observed growth arrest in contrast to the bactericidal phenotype resulting from depletion of *lpxC*, which is involved in LPS biosynthesis (Figure S2C). In other cases, we simply observed slower growth, as in the case of *lptA* (involved in LPS transport) and *parC* (a subunit of topoisomerase). The contrast between *lpxC* and *lptA* phenotypes, both involved in LPS synthesis and transport, potentially suggests that *lpxC* may be the more vulnerable, more attractive target in the LPS pathway. Moreover, even within the same complex, we observed widely disparate growth phenotypes with similar levels of depletion, suggesting differential vulnerabilities even within the same complex, as previously observed in Mycobacteria within the DNA replication-related enzyme complexes.^17^ For example, we observed variable growth defects when we knocked down the terminal components of the essential LPS transport complex, which is composed of LptD and LptE, which together form the outer membrane transport channel, and LptA, which forms the periplasmic bridge through which LPS is transferred before the LptD/E complex delivers it to the outer surface. Despite similar levels of target depletion in *lptD* and *lptA* (log2 fold change (L2FC) of-4.27 and-3.67, respectively, relative to induced control), the degree of growth defect was more pronounced in *lptA*-CRISPRi^R^ than in *lptD*-CRISPRi^R^ (Figure S2B and S2D). Further, despite knockdown of L2FC of-3, *lptE*- CRISPRi^R^ had no growth defect at all, phenocopying what has been observed in an arabinose-inducible *lptE* knockdown strain^33^ (Figure S2B and S2D). Similarly, while depletion of both *gyrA* and *gyrB* (L2FC of-4.4, and -2.9, respectively) of the DNA topology gyrase enzyme had minimal impact on growth, depletion of the *parC* subunit (L2FC of-3.4) of the related topoisomerase IV enzyme, led to a dramatic growth defect (Figure S2B and S2D). Thus, perturbing essential genes across similar processes (cell envelope modulating *murA* and *lpxC*), within the same pathway (*lptA*, *lptD*, *lptE*), or having similar functions (*gyrA* and *parC*) led to diverse growth outcomes, which implies differential target vulnerability that has important implications for drug discovery efforts.^17^

### CRISPRi to probe mechanism of action of small molecule inhibitors

We used the tightly regulated CRISPRi^R^ system first to create partial knockdown mutants to identify chemical genetic interactions between specific genes with anti-pseudomonal molecules, and then to generate maximal knockdown mutants to measure specific transcriptional signatures in response to pronounced essential target depletion. We first leveraged the tunability of the rhamnose CRISPRi system to examine chemical genetic interactions between the depletion mutants and small molecule inhibitors.^34–37^ We used sub-lethal rhamnose doses to achieve partial gene depletion of the 16 selected essential genes and measured the activity of 13 known small molecules targeting DNA replication, transcription, protein translation, and cell envelope synthesis in these strains using standard growth-based assays (Figure S4). We observed sensitization of a CRISPRi^R^ mutant to a chemical inhibitor in 12 of 13 cases (Figure S5). For example, knockdown of *gyrB*, *folA*, and *dxr* (involved in isoprenoid biosynthesis) sensitized to their respective inhibitors, novobiocin, trimethoprim, and fosmidomycin (Figure 4A). Of note, knockdown of the other gyrase subunit, *gyrA*, also caused novobiocin sensitization, even though it does not bind GyrA directly, revealing a functional interaction between the two subunits in the target protein complex. None of the DNA-synthesis-related CRISPRi^R^ strains showed shifts in susceptibility to ciprofloxacin, which directly binds GyrA and ParC in complex with DNA to stabilize DNA breaks (Figure 4A).^38^

**Figure 4.**
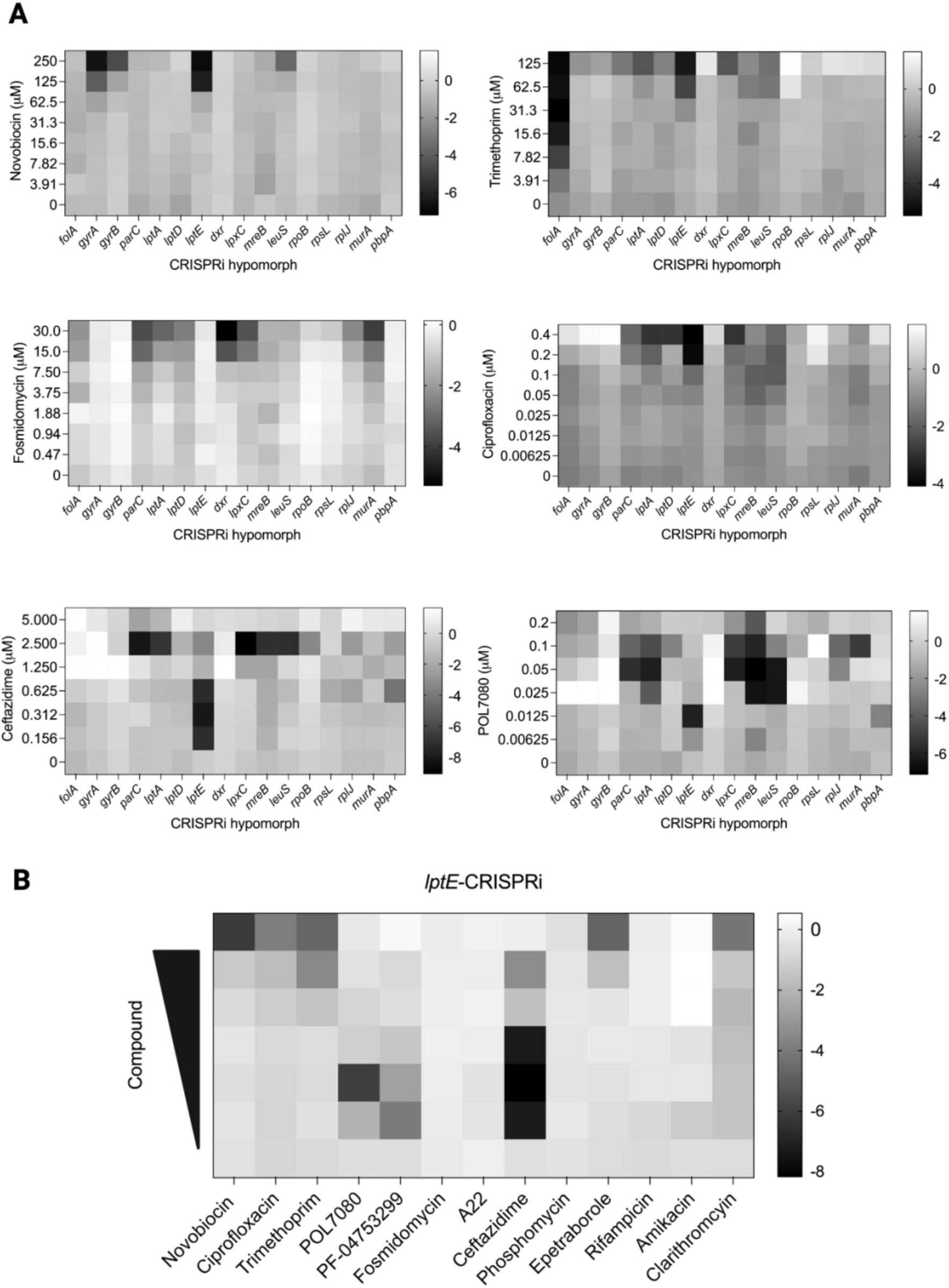
CRISPRi^R^ genetic perturbation leads to altered chemical sensitivities. (A) Heat map of the log_2_ fold change (L2FC) in growth of CRISPRi^R^ strains relative to *control*-CRISPRi^R^ strain. CRISPRi^R^ strains were grown in induced conditions albeit with sub-lethal concentrations of rhamnose. (0.05% rhamnose: *folA-*, *gyrA-*, *gyrB-*, *lptE-*, *rpoB-*, and *pbpA*-CRISPRi^R^ strains; 0.025% rhamnose: *lptA-*, *lptD-*, *dxr-*, *mreB-*, *leuS-*, *rpsL*, and *rplJ*-CRISPRi^R^; 0.0125% rhamnose: *lpxC-* and *murA*-CRISPRi^R^.) Growth (OD_600_) was measured for each strain after 24 hours in the presence of chemical inhibitor and L2FC of growth relative to the growth of the *control*-CRISPRi^R^ strain was calculated. CRISPRi^R^ strains displayed variable sensitivities to chemical inhibitors. (B) Heat map of the L2FC in growth of the *lptE*-CRISPRi^R^ strain relative to *control*-CRISPRi^R^ strain reveals broad sensitization to a number of chemical inhibitors with different MOA. Scale indicates L2FC in growth relative to *control*-CRISPRi^R^ strain.

Meanwhile, in the case of cell wall active agents, we often observed chemical genetic interactions linking the cell wall and membrane integrity. For example, the β-lactam ceftazidime was not only more potent against the canonical penicillin-binding protein strain, *pbpA*-CRISPRi^R^, relative to wild-type, but also against membrane integrity depleted strains such as *lpxC*- (involved in LPS biosynthesis), *lptA*-, *lptE*-, and *mreB*-CRISPRi^R^. Similarly, the cyclic peptide LptD inhibitor, POL7080, not only had enhanced potency towards its target LPS pathway-specific strains such as *lptA*-, *lptE*-, *lpxC*-CRISPRi^R^, but also towards peptidoglycan synthesis-linked *mreB*-CRISPRi^R^ (Figure 4A). Interestingly, POL7080 also had an interaction with the protein synthesis-related *leuS*-CRISPRi strain, potentially linking mistranslation to envelope stress^39^. Alternatively, this observed interaction could be due to operon structure with *lptE* being immediately downstream of *leuS*, resulting in *lptE* depletion in the *leuS*-CRISPRi^R^ strain (*lptE* L2FC of-2.6 in the *leuS*- CRISPRi strain, Table S1). Notably, although the *lptE*-CRISPRi^R^ strain was not growth impaired, it was broadly sensitized across the drug panel indicating a vital role for LptE in maintaining the cell envelope barrier in *P. aeruginosa* (Figure 4B). Thus, growth inhibitor sensitization studies with CRISPRi-based hypomorphs showed direct as well as more complex chemical genetic interactions, which can provide insights into gene/pathway functions in *P. aeruginosa*.

### Expression profiling on CRISPRi strains sheds light on *P. aeruginosa* response to genetic perturbations

We also leveraged the rhamnose-based system to achieve maximal knockdown of essential genes in *P. aeruginosa* in order to measure genome-wide transcriptional responses to specific genetic depletions. These target specific transcriptional changes were then mapped to the equivalent changes due to chemical perturbation of the corresponding essential pathways, utilizing our previously reported PerSpecTM method.^40^ We initially performed transcriptomic analysis of the 16 CRISPRi^R^ knockdown strains after 6 hours of induction with maximum concentrations of rhamnose, as previously performed using the arabinose CRISPRi^A^ system. However, we noted that the degree of transcriptional change in induced strains relative to *control*-CRISPRi strains was less for rhamnose-based strains than for arabinose ones at this same time point (Figure 2E). We thought this could be due to the relatively shorter induction period of the rhamnose system, taking into account that the leak in the arabinose system would have provided longer, sustained activation of the system (Figure 2E). When we extended the period of rhamnose induction to 10 hours, we measured similar levels of target gene knockdown as at 6 hours but were able to capture broader genome-wide responses to perturbation, based on the number of differentially expressed genes (DEGs) (Figure S6A and S6B).

Previously, PerSpecTM analysis of a large *P. aeruginosa* transcriptomic dataset accurately predicted the mechanism of action (MOAs) of known and novel small molecule inhibitors.^40^ Predictions were made based on aligning inhibitor-induced bacterial transcriptional responses with reference profiles from known-MOA compounds and from arabinose-inducible target knockdowns. Compounds sharing a mechanism showed correlated transcriptional signatures. Notably, the transcriptional response to a small molecule could also be correlated with the profile resulting from genetic knockdown of its cognate target. Here, using CRISPRi^R^ strains with the rhamnose-inducible knockdown after 10 hours of induction, we observed stronger Pearson correlations between gene knockdown profiles and their corresponding small molecule treatments than with profiles generated using the arabinose-based system for 15 of 25 chemical perturbations (Figure 5A and Figure S7). Moreover, matching transcriptional profiles between small molecules and their CRISPRi-depleted target strains proved more sensitive for identifying target relationships than mapping small molecules to target genes based on CRISPRi hypersensitivity growth phenotypes. For example, while strains depleted of *rpoB*, *rpsL*, and *rplJ* (which encode subunits of RNA polymerase, the 30S ribosome, and 50S ribosome, respectively) were not hypersensitized to transcription or translation inhibitors (rifampicin, tetracyclines, or macrolides) in a growth-based MIC_90_ assay (Figure S5), a robust interaction emerged based on correlation of transcriptional profiles for *rpoB*-CRISPRi with rifampicin treatment and *rpsL*-CRISPRi and *rplJ*-CRISPRi strains with treatment by two protein synthesis inhibitor classes (Figure 5A). Similarly, while we saw no sensitization of any of the DNA synthesis-associated CRISPRi^R^ strains to ciprofloxacin (Figure S5), the transcriptional responses to fluoroquinolones and cisplatin were significantly correlated with the response to *parC* knockdown (Figure 5A). These observed correlations demonstrate the greater sensitivity of a gene expression readout relative to an OD_600_-based growth measurement for identifying chemical genetic interactions.

**Figure 5.**
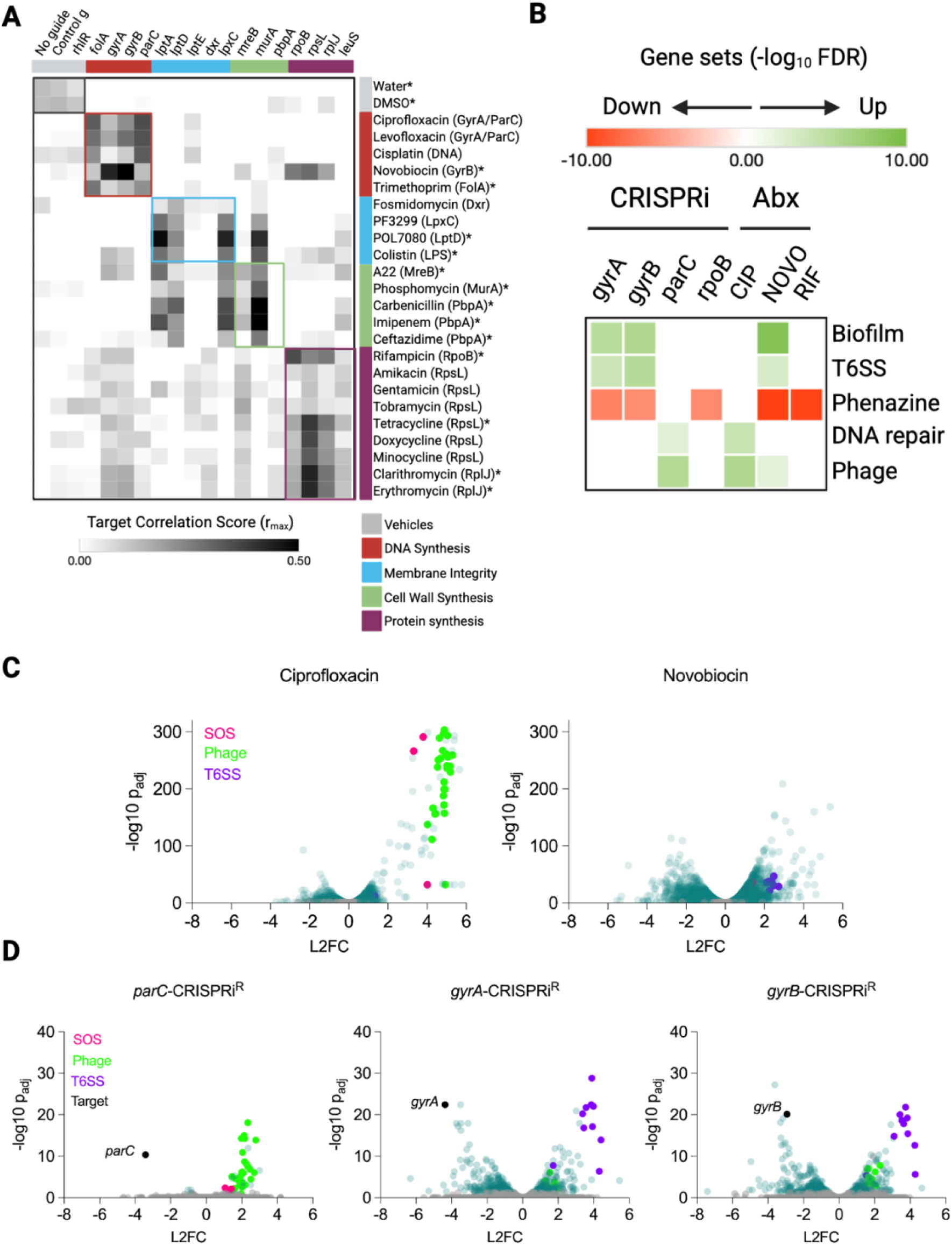
Transcription profiling of CRISPRi^R^ knockdown strains reveals chemical genetic interactions. (A) PerSpecTM analysis of chemical inhibitors queried against CRISPRi^R^ strains. Pearson correlations ( r̅_max_ ) of z-scored expression profiles from 90-min chemical inhibitor treatment on PA14 with 10h of gene depletion of CRISPRi^R^ strains are shown as a heat map. Responses to all chemical inhibitors were characterized in PA14 except fosmidomycin, which was characterized in a *dxr*-hypomorph strain due to the lack of wild-type activity. CRISPRi strains and chemical inhibitors with the same or related mechanism of action (MOA) are color coordinated. The transcription profiles of chemical inhibitors largely cluster according to their MOA. Cell envelope genetic perturbations show a degree of crosstalk between membrane integrity and peptidoglycan synthesis chemical inhibitors. Asterisk indicates higher Pearson correlations between transcription profiles of a chemical inhibitor and genetic perturbation of its predicted target with CRISPRi^R^ than with CRISPRi^A^(Figure S7).^40^ (B) Heat map showing gene sets enriched after CRISPRi^R^ genetic perturbation relative to the control-CRISPRi^R^ strain or after chemical perturbation relative to vehicle control. Enrichment is quantified by the false discovery rate measured using STRING-DB with green and red colors indicating upregulated and downregulated gene sets, respectively. White boxes indicate no functional enrichment of a gene set. CRISPRi^R^ perturbation revealed a distinct bacterial response to gyrase versus topoisomerase IV depletion. The DNA damage response (SOS) leading to upregulation of DNA repair and phage assembly genes, which is a hallmark of fluoroquinolone exposure, was observed after topoisomerase IV knockdown (*parC*-CRISPRi^R^) but not gyrase (*gyrA*-, *gyrB-*CRISPRi^R^) nor RNA-polymerase (*rpoB*-CRISPRi^R^) knockdown. (C) Volcano plots showing gene expression in PA14 in response to ciprofloxacin and novobiocin treatment. Ciprofloxacin induced genes involved in SOS response (pink) and phage assembly (blue). (D) Volcano plots showing gene expression in response to CRISPRi^R^ knockdown of topoisomerase genes. *parC* knockdown elicited upregulation of SOS response and phage assembly genes whereas *gyrA* and *gyrB* knockdown led to type VI secretion system (T6SS) upregulation.

While molecule-molecule, molecule-gene, or gene-gene correlations were based on the genome-wide patterns of transcriptional response, agnostic to specific gene identities, the actual genes and pathways whose expression is perturbed by a molecule or gene knockdown can provide insight into a molecule’s MOA or a gene’s function. We performed differential gene expression analysis followed by functional enrichment using STRING-DB to identify pathways that were differentially induced in response to specific perturbations (Figure 5B). Interestingly, the SOS response—a hallmark DNA damage and repair— was induced when *parC* but not *gyrA* or *gyrB* were depleted, suggesting that reduction of ParC’s decatenation activity results in accumulation of single stranded DNA breaks, double stranded DNA breaks, or unresolved replication intermediates that trigger the RecA-mediated SOS response. Key SOS genes such as *recA*, *recN*, and phage-related genes downstream of the SOS response were among the most highly upregulated upon *parC* depletion or from ciprofloxacin treatment but not novobiocin treatment (Figure 5C and 5D). In contrast, the loss of gyrase’s negative supercoiling activity alone in the *gyrA* and *gyrB* depleted strains, per se, did not cause sufficient DNA damage to induce the SOS response (Figure 5D). Further, in that fluoroquinolones are highly activating of the SOS response, these results are consistent with what has now been long established, that fluoroquinolones do not act simply by inhibiting enzymatic function at least in the case of gyrase; instead, they act by a poisoning mechanism.^38,41^ This result is in contrast to novobiocin, which is not only more potent against the *gyrB* and *gyrA* depleted strains (Figure 4A), but also phenocopied the transcriptional consequences of *gyrB* and *gyrA* depletion (Figure 5A). These results together point to a simple enzymatic inhibition mechanism for novobiocin.

Together, these findings show that CRISPRi-mediated knockdown often mimics the transcriptional effects of chemical inhibition of essential pathways. Moreover, some knockdowns reveal unique response signatures, helping to link specific gene functions to stress responses in *P. aeruginosa*.

## DISCUSSION

We present an improved CRISPRi system using the *S. pasteurianus* dCas9 for *P. aeruginosa* with enhanced control over gene knockdown, enabling robust genetic perturbation studies of essential genes. In contrast to a recently reported CRISPRi doxycycline-induced *S. pyogenes* dCas9 system for *P. aeruginosa*^14^, we focused on small hydrophilic, sugar-based induction systems as they are less susceptible to efflux pumps thereby increasing the likelihood that they could be applied broadly to *P. aeruginosa* strains including clinical isolates which generally have high baseline efflux pump activity, and might show less mutant to mutant variability in inducer cell entry and intracellular accumulation. Building on a prior system utilizing *S. pasteurianus* dCas9 (dCas9_Spa_), we optimized expression control to target essential genes not only to ensure robust target repression, but also importantly, to ensure no activation in the absence of inducer to ensure stable maintenance of individual mutants or pools of mutants with propagation. This feature is critical for enabling studies of essential gene function. In contrast to other reported sugar-based systems relying on arabinose-induction (P_araBAD_) or IPTG-induction (P_lac/tac_) of dCas9 with constitutive promoters driving sgRNA expression^13,15^, which resulted in leaky expression and unintended gene repression in the absence of inducer, we employed a different approach. We used the rhamnose-inducible P_rhaBAD_ promoter to regulate both *dcas9* and sgRNA expression, inspired by the tightly regulated CRISPRi system in *Mycobacterium spp*.^16^ To accommodate the larger P_rhaBAD_ promoter, we divided the CRISPRi system across two integrating vectors: one expressing dCas9 for integration into the *ϕ CTX* attachment site, thereby generating a parental CRISPRi strain (PA14::*dcas9*), and the other expressing sgRNAs for integration into the *att*Tn7 site (Figure 2A). This modular design enabled efficient cloning and integration via conjugation of compact sgRNA-carrying miniTn7 plasmids into PA14::*dcas9*, unlike the inefficient integration of a single, large dual-function plasmid (Figure 1A). The combination of tight regulation and high conjugation efficiency facilitated rapid and facile—even pooled—construction of CRISPRi knockdown libraries in *P. aeruginosa* (Figure S3B). We also identified a new PAM sequence, NNGTAT, which in combination with the three known PAMs will expand the scope of guide locations in the target genome, further enabling genome-wide perturbation.

Using our dCas9_Spa_-based system, we knocked down 16 essential genes involved in diverse cellular processes, including DNA replication, transcription, translation, and envelope biogenesis. As expected, we observed highly variable phenotypes depending on the gene and the degree of knockdown that could be achieved, based on the vulnerability of each gene *i.e.*, its ability to tolerate depletion before impacting phenotype. For instance, *lpxC* depletion which impacts LPS synthesis was bactericidal, whereas *murA* knockdown which impacts peptidoglycan synthesis was bacteriostatic, highlighting differential phenotypes across envelope biogenesis pathways. Moreover, within the same complex, similar levels of depletion of different components resulted in widely disparate growth phenotypes, as previously described in Mycobacteria^17^ and here exemplified in depletion of Lpt complex components *lptD*, *lptA*, and *lptE* (Figure S2D). Assuming the relative depletion of their mRNA levels translates to protein depletion, these differences suggest that LptA may be the most vulnerable target in the pathway and might be the stoichiometric bottleneck for Lpt complex formation and function.^42,43^ Similarly, amongst DNA topology modulating type II topoisomerase enzymes, *parC* depletion caused severe growth defects, while depletion of gyrase genes, *gyrA* and *gyrB*, had only modest effects (Figure S2D).

For a given gene, we typically observed changes in the transcription profile occurring even with low levels of depletion as the cell may sense stress or imbalance and attempt to compensate. With greater depletion, increased sensitivity to inhibitors of the target became manifest because of less available functional target. Only with extreme levels of depletion where there is no longer sufficient functional target for normal viability did we observe a detectable impact on survival and growth.^44,45^ Thus, changes in transcriptional programs, followed by MIC_90_ shifts, and then finally growth defects formed a typical hierarchy of sensitivity to target depletion. This hierarchy was demonstrated by a number of genes in this study. For example, in the case of *folA* and *mreB*, the corresponding uninduced arabinose-based strains had no growth defects but did display shifts in MIC_90_ to cognate inhibitors or morphologic changes, reflecting more sensitive detection of depleted function. Further, while strains depleted of *rpoB*, *rpsL*, *rplJ*, and *parC* were not hypersensitized to cognate inhibitors of the corresponding gene product or pathway in a growth-based MIC_90_ assay, the transcriptional profiles resulting from genetic depletion correlated well with treatment of wild-type PA14 by their corresponding cognate inhibitor.

Comparisons of these phenotypes in different genetically depleted strains can reveal novel aspects of gene function, particularly when discordance is detected among similar targets, such as those within the same complex. For example, while *lptE* depletion did not impair growth, knockdown sensitized (4 to 8-fold shift in MIC_90_) to several anti-pseudomonal molecules including POL7080 and PF-04753299, which target LPS synthesis; novobiocin and clarithromycin, which are typically excluded by the *P. aeruginosa* outer membrane; and ceftazidime, the β-lactam peptidoglycan inhibitor. We also observed moderate sensitization (2 to 4-fold shift in MIC_90_) to ciprofloxacin, trimethoprim, and epetraborole. These results suggest that despite LptE’s relative invulnerability to depletion with respect to growth impairment, it potentially plays a unique role in outer membrane integrity, relative to other, more growth vulnerable Lpt complex proteins such as LptA, which may serve as the stoichiometric determining subunit of the Lpt complex.^46–48^

Similarly, *parC* knockdown differed from *gyrA* and *gyrB* depletion, with the former inducing the SOS response including DNA repair and phage assembly pathways, which is the hallmark bacterial response to DNA breaks, thereby phenocopying the response to fluoroquinolone exposure. In contrast, a bit surprisingly, the gyrase depletion of both *gyrA* and *gyrB* subunits resulted in the strong upregulation of biofilm genes and the type VI secretion system and repression of quorum sensing genes in the PQS pathway, without activation of the SOS response.^49^ The mechanism underlying this surprising relationship between gyrase depletion and these virulence mechanisms remains to be elucidated. Nevertheless, these divergent transcriptional responses to depletion underscore fundamental differences in the mechanisms by which *P. aeruginosa* senses and responds to disruption of gyrase, topoisomerase IV, and/or their regulatory pathways. Further, they highlight the complexity of fluoroquinolone activity in contrast to novobiocin activity, with the latter acting as a direct inhibitor of GyrB enzymatic function while fluoroquinolones act via a topoisomerase poisoning mechanism. Thus, genetic perturbation can result in phenotypic consequences distinct from growth to reveal new insight into gene function or small molecule MOA.^40^

While the application of CRISPRi to this set of 16 genes provided insight into some chemical interactions and target vulnerabilities, this study nevertheless was limited in scope. Phenotypic outcomes are dependent on the degree of knockdown achieved and the functional reserve of a target *i.e.,* ability of the cell to tolerate target depletion. In the case of CRISPRi, the nature of the selected sgRNA, including PAM strength, guide location, and guide length, is an important factor that determines target knockdown levels and consequently, cellular responses. The other important factor determining phenotypic outcomes is the inherent vulnerability of the target gene. Although we attempted to select the strongest PAM for each gene in order to elicit the most dramatic phenotypes approximating complete target depletion, based on the mKate2 reporter assay and guide position closest to the transcription start site, it is possible that we did not in reality achieve this goal in all cases. Thus, suboptimal guides resulting in insufficient or variable target depletion could explain some of the differential growth and inhibitor susceptibility profiling phenotypes that we observed. To identify the strongest sgRNA in an unbiased manner, a more comprehensive pooled CRISPRi knockdown library, which includes multiple sgRNAs across a range of strengths, would need to be systematically evaluated.^17^ A related caveat in assessing the level of target knockdown by CRISPRi is that mRNA knockdown levels do not always accurately reflect the extent of protein depletion; this distinction is important to consider when interpreting the relationship between genetic vulnerability and cellular phenotype.

Another limitation of CRISPRi is that it can generate polar effects in bacterial systems, including in *P. aeruginosa*, due to the operonic organization of bacterial genomes. Thus, the targeted knockdown of one gene could also impact genes downstream in the operon leading to effects on observed phenotypes that are not due to the specific CRISPRi targeted gene. We observed such effects with some of our targets, like *mreB*, wherein we observed downregulation of downstream *mreC* and *mreD* in *mreB*-CRISPRi^R^, and potentially *lptE* in *leuS*-CRISPRi^R^, which lies downstream of *leuS*. However, clearly this is not a universal phenomenon, since in other CRISPRi strains, such as *lptA-*CRISPRi^R^, with the *lptB* gene located downstream, we did not observe any polar effects. Therefore, large scale CRISPRi-based functional or phenotypic characterization needs to include controls that account for these polar effects. Despite all of the caveats in using CRISPRi to study the consequences of genetic depletion, we nevertheless found that the observed phenotypes consistently aligned with expected target depletion effects.

In conclusion, the rhamnose-inducible CRISPRi system enables precise and tunable repression of essential genes in *P. aeruginosa*, thus making it well-suited both for detailed analysis of individual essential genes and for large-scale, genome-wide functional genomic studies and chemical-genetic screens— approaches that have already proven powerful in other bacterial pathogens such as *E. coli*^50–54^, *B. subtilis*^11,55^, mycobacteria^17,56^, and most recently *P. aeruginosa*^14^. Notably, this system supports the generation and stable maintenance of comprehensive pooled mutant libraries inclusive of essential genes, agnostic of strain efflux pump background, thereby allowing for comprehensive, systematic assessment of individual gene vulnerabilities across a large number of clinical isolates. Such studies will provide critical insights for target prioritization in antibiotic discovery efforts and will help to accelerate the development of new therapeutics against *P. aeruginosa*, a pathogen that has proven historically recalcitrant to traditional antibiotic discovery strategies.

## METHODS

### CRISPRi cloning and strain construction

All sgRNA sequences and primer information are shown in Table S1.

#### First generation P_araBAD_ CRISPRi^A^ system

The catalytically inactive Cas9 (dCas9) from *Streptococcus pasteurianus* and *Streptococcus pyogenes* were purchased from Addgene (Watertown, MA)^13,15^; the dCas9 from *Streptococcus thermophilus* was gifted by Jeremy Rock^16^. The *dcas9* gene was cloned into a modified pUC18T-miniTn7T-pBAD-Gm vector previously described by our group under the control of the AraC-araBAD promoter system using standard 2-piece Gibson Assembly.^57^ Three BsmBI restriction sites were removed by QuikChange XL mutagenesis kit (Agilent, Santa Clara, CA). A custom gene block was synthesized containing a 5’ HindIII restriction site, *E. coli* ProD promoter, two unique BsmBI restriction sites position the sgRNA at the transcriptional start site of ProD promoter, rrnB T1 terminator, and a KpnI restriction site. The gene block and pUC18T-miniTn7T-pBAD-dcas9-Gm plasmid were restriction digested with HindIII and KpnI, and ligated together (T4 DNA ligase, New England Biolabs, Ipswich MA) to produce the first generation CRISPRi^A^ plasmid pUC18T-miniTn7T-pBAD-dcas9-pD-Gm (9326bp). sgRNAs were cloned into the CRISPRi^A^ plasmid using standard CRISPRi protocols established in mycobacteria using BsmBI cut sites.^16^ Another iteration of the CRISPRi^R^ dCas9_Spa_ was constructed on a pUC18-miniTn7T-pRha- dcas9-pD-Gm vector under the control of the RhaRS-rhaBAD promoter system (10140bp). All constructs were transformed into 10-beta *E. coli* (New England Biolabs, Ipswich MA). Following rapid plasmid sequencing, tetrapartite bacterial conjugation was carried out in a 1:1:1:1 mixture of recipient (PA14), 10- beta *E. coli* donor cells harboring the construct of interest, and *E.coli* cells harboring the helper plasmids pRK2013 and pTNS2^24^. Recipient PA14 cells were incubated at 42 °C for 2 hours prior to the tetrapartite mating. Cells were mated at 37 °C for 24 hours on cellulose membranes, resuspended in 400 μL of LB and plated on LB agar containing 15 μg/mL gentamicin and 5 μg/mL irgasan. Colonies were screened for correct integration of the miniTn7 plasmid using primers 10 and 12.

#### Constitutive mkate2 strain used with first generation CRISPRi^A^

The fluorescent gene *mkate2* was custom synthesized as a gene block under the control of a constitutive P_trc_ promoter. Gene blocks of *mkate2* with mutated PAM sites were also synthesized. The ptrc-mkate2 fragment was PCR amplified using primers 1 and 2 and double digested with SpeI and SacI. The digested fragment was ligated into double digested miniCTX1^58^ plasmid and transformed into 10-beta *E. coli*. Following rapid plasmid sequencing, tripartite bacterial conjugation was carried out in a 1:2:2 mixture of recipient (PA14), 10-beta *E. coli* donor cells harboring the CTX-*mkate2* construct, and *E.coli* cells harboring the helper plasmid pRK2013. Recipient PA14 cells were incubated at 42 °C for 2 hours prior to the tripartite mating. Cells were mated at 37 °C for 24 hours on cellulose membranes, resuspended in 400 μL of LB and plated on LB agar containing 15 μg/mL gentamicin and 5 μg/mL irgasan.

#### Second generation P_rhaBAD_ CRISPRi^R^ system

the second generation system split the CRISPRi components into two plasmids; moving the *dcas9* gene with a rhamnose-inducible promoter on a miniCTX1 vector to create a PA14:dcas9 strain that can be used as the recipient strain for any guide, and moving the sgRNA with a rhamnose-inducible promoter on the smaller pUC18-miniTn7T-Gm plasmid (RhaBAD vector: 6470bp) to facilitate cloning and conjugations. The miniCTX1-ptrc-mkate2-tet (see constitutive mkate strain above) and the pUC18-miniTn7T-pRha-dcas9-pD-Gm vectors were double digested with HindIII and SacI to excise the ptrc-mkate fragment and RhaRS-rhaBAD-dcas9 fragment, respectively. Linearized miniCTX1 and the RhaRS-rhaBAD-dcas9 fragment were ligated together to create a miniCTX1-pRha-dcas9-Tet plasmid (11874bp). Following transformation into 10-beta *E.coli* and rapid plasmid sequencing, the plasmid was conjugated into PA14 using the tripartite mating strategy. Positive conjugants were selected on 100 μg/mL tetracycline and 5 μg/mL irgasan and screened using primers 5 and 6 to check for correct CTX integration^58^ at the *glmS* site and primers 3 and 4 to check for the presence of *dcas9*. To remove the tetracycline resistance cassette in the FRT region of the integrated miniCTX1 plasmid, a clone of PA14:CTX- dcas9 was mixed in a 1:5 ratio with *E.coli* cells carrying the SM10-pFLP2 flippase plasmid and incubated at 37 °C for 24 hours on cellulose membranes. Conjugants were selected on LB agar 5 μg/mL irgasan. After overnight incubation at 37 °C, colonies were patched onto LB agar with 5 μg/mL irgasan, and LB agar with 100 μg/mL tetracycline + 5 μg/mL irgasan. After overnight incubation at 37 °C, colonies that grew on LB 5 μg/mL irgasan and not 100 μg/mL tetracycline were streaked onto LB agar with 15% sucrose to remove the SM10-pFLP2 plasmid. Single colonies that grew overnight at 37 °C were patched onto LB 100 μg/mL carbenicillin and LB 100 μg/mL tetracycline to check for the loss of the SM10-pFLP2 plasmid and loss of the tetracycline cassette in the FRT region. Positive clones were screened by PCR to confirm the absence of the FRT region and presence of RhaBAD-dcas9 using primers 5 and 7, and 3 and 4, respectively.

#### Second generation sgRNA plasmid

a Tn7 vector with a rhamnose-inducible promoter controlling sgRNA expression was constructed to facilitate the cloning of sgRNAs into the parental PA14::*dcas9* strain. Q5 mutagenesis was performed on pUC18-miniTn7T-pRhabad-dcas9-pD-Gm plasmid to remove the *dcas9* coding sequence and ProD promoter, and bring the Rhabad promoter transcriptional start site in line with the sgRNA generating pUC18-miniTn7T-pRhadbad-Gm. Constructs were transformed into 10-beta *E. coli* and rapid plasmid sequenced. sgRNAs were cloned into the CRISPRi^R^ plasmid using standard CRISPRi protocols established in mycobacteria using BsmBI cut sites.^16^ Sequence-confirmed sgRNA plasmids were conjugated into PA14:CTX-dcas9 (PA14::*dcas9*) parental strain using the tetrapartite method described above. Positive conjugants were selected on LB agar with 15 μg/mL gentamicin and 5 μg/mL irgasan and PCR screened for correct *attB*-Tn7 integration using primers 10 and 12.

#### Constitutive mkate2 strain used with second generation CRISPRi^R^

The miniCTX-ptrc-mkate plasmid was double digested with BamHI and SacI to remove the ptrc-mkate fragment. The recipient sgRNA vector pUC18-Tn7T-pRhabad-Gm was also double digested and ligated with the ptrc-mkate fragment. Constructs were used to clone in *mkate2* guides using BsmBI cut sites and conjugated into PA14::*dcas9* parental strain to test the efficacy of the P_rhaBAD_ CRISPRi^R^ system.

### dCas9 Western blot

Overnight cultures grown in LB broth (37 °C, 250rpm) were diluted 100-fold in fresh LB with or without arabinose (0.5%) and grown until OD_600_ reached mid log. Twenty mL of cells were pelleted and resuspended in 1 mL of lysis buffer (50 mM Tris pH 7.5, 1 mM EDTA, 150 mM NaCl, 5 μg/mL DNase I, 0.5 mg/mL lysozyme, Roche protease inhibitor cocktail). Cells were lysed by sonication and supernatants clarified by centrifugation at 13,000g at 4 °C for 10 minutes. Protein concentrations were quantified using the micro-BCA assay. Protein extracts (20 μg) were separated by SDS-PAGE (4-20% Biorad TGX gels) and transferred to PVDF membrane. Membranes were incubated overnight with primary anti-6×his mouse (Sigma #SAB2702218) in a 1:5,000 dilution or anti-RpoA mouse (Sigma #F3165) in a 1:20,000 dilution at 4 °C. Membranes were washed and incubated with anti-mouse HRP at 1:10,000 dilution for one hour at room temperature.

### mKate2 fluorescence knockdown

Overnight cultures grown in LB broth (37 °C, 250rpm) were diluted 100-fold in M9 minimal media (M9 salts, 0.4% glucose, 2 mM MgSO_4_, 0.1 mM CaCl_2_) and grown until OD_600_ reached mid log. Cultures were diluted to OD_600_ 0.01 and added to a 96-well plate (Nunc 96-well black, optical bottom plate, Thermo Fisher Scientific, Waltham MA) containing the appropriate amount of arabinose/rhamnose. The plate was incubated in a humidity chamber with shaking at 37 °C in a Spark Multimode Reader (Tecan, Männedorf Switzerland) where fluorescence (Ex 588nm, Em 633nm) and OD_600_ were measured every hour for 24 hours. PA14 was used to subtract background fluorescence. First generation CRISPRi^A^ *mkate2* knockdown strains were compared to a control-CRISPRi^A^ strain that carried a scrambled control sgRNA that had no impact on fluorescence or growth in minimal media. Second generation CRISPRi^R^ *mkate2* knockdown strains were compared to a no sgRNA control strain that carried a guide-less pUC18-Tn7T-ptrc-mkate- pRhabad-Gm plasmid.

### CRISPRi growth kinetics & Minimum inhibitory concentration measurements

#### Growth in liquid

Overnight cultures grown in LB broth (37 °C, 250rpm) were diluted 100-fold in fresh LB broth or with LB- arabinose or rhamnose to pre-activate the system and grown until OD_600_ reached mid log. Cultures were diluted to OD_600_ 0.001 and 25 μL added to a 384-well plate (Nunc 384-well clear polystyrene plates, Thermo Fisher Scientific, Waltham MA) containing the appropriate amount of 2X arabinose/rhamnose or chemical inhibitor. The plate was incubated in a humidity chamber with shaking at 37 °C in a Spark Multimode Reader (Tecan, Männedorf Switzerland) where OD_600_ was measured every hour for 24 hours. Chemical genetic interaction profiles were calculated by the fold change in growth of the *target*-CRISPRi^R^ strain relative to the *control*-CRISPRi^R^ strain.

For kill kinetics experiments, cultures were diluted to OD_600_ 0.0001 in 15 mL glass tubes containing the appropriate amount of rhamnose. The tubes were incubated at 37 °C 250rpm for up to 12 hours. One hundred μL aliquots were collected, 10-fold serially diluted in 1X PBS, and plated on LB agar in 10 μL spots. Plates were incubated overnight at 37 °C in a humidity box. Colonies were counted and plotted as CFU/mL.

#### Growth on solid media

Overnight cultures grown in LB broth (37 °C, 250rpm) were diluted 100-fold in fresh LB broth and grown until OD_600_ reached mid log. One hundred μL aliquots were collected, 10-fold serially diluted in 1X PBS, and plated on LB agar with the appropriate amount of arabinose in 10 μL spots. Plates were incubated overnight at 37 °C in a humidity box.

### Microscopy

Overnight cultures grown in LB broth (37 °C, 250rpm) were diluted 100-fold in fresh LB broth and grown until OD_600_ reached mid log. Cultures were diluted to OD_600_ 0.001 and 75 μL added to a 96-well plate with 75 μL of the corresponding 2X rhamnose or arabinose and incubated for 19 hours. One hundred μL of cells were collected and mixed with 100 μL 1X PBS, and spun down at 9000 rcf for 10 minutes. The supernatant was removed and the pellet suspended in 50 μL 1X PBS with 2.5 μg/mL of FM-143x (Invitrogen #F35355). Samples were incubated on ice for 15 minutes. One hundred and fifty μL of 1X PBS were added on top, pelleted and resuspended in 50 μL 4% paraformaldehyde in 1X PBS. Samples were incubated on ice for 20 minutes. One hundred and fifty μL 1X PBS were added on top, pelleted, and resuspended in 20 μL 1X PBS. One μL of stained cells and 3 μL of ProLong™ Gold Antifade reagent (Invitrogen #P36930) were added to a glass microscope slide, covered with a coverslip and imaged on a Zeiss confocal microscope after 24 hours of drying.

### RNA-Seq experiments

RNA-Seq experiments were carried out in biological triplicate in 384-well microplates (Nunc 384-well clear polystyrene plates, Thermo Fisher Scientific, Waltham MA). Overnight cultures of CRISPRi^R^ strains grown in LB broth (37 °C, 250rpm) were diluted 50-fold in fresh LB in a 96 deep-well block. Bacterial cultures were grown with shaking (37 °C, 250r pm) to mid-log phase, and 30 μL of cultures were mixed with equal volume of 2X rhamnose working solution, yielding final conditions containing 1x10^8^ CFU/mL bacteria and with 0%, 0.0125%, or 0.05% rhamnose in LB. CRISPRi^R^ strains were treated at 37 °C without shaking in a humidity chamber for 6 hours. After incubation, 50 μL of cultures were mixed with 25 μL 3X RNAgem Blue Buffer (Zygem, Charlottesville VA), and chemically lysed by incubation at 75 °C for 10 minutes in a thermocycler. Total RNA was extracted and purified using the Direct-zol RNA Miniprep Plus kit and TRIzol reagent (Zymo Research, Irvin CA). RNA quality and quantity were analyzed using the RNA ScreenTape with 2200 TapeStation (Agilent, Santa Clara CA).

An alternative experimental setup was used for the kinetics experiment. After diluting the overnight cultures 50-fold in fresh LB in a 96-well block, cells were back diluted to OD_600nm_ ∼ 0.05 in 96-well blocks with/without 0.05% rhamnose. Cells were grown until OD_600nm_ 0.4-0.6 was reached, 120 μL of culture was mixed with 3X RNAgem Blue Buffer (Zygem, Charlottesville VA), and chemically lysed by incubation at 75 °C for 10 minutes in a thermocycler. Remaining cultures were back diluted to OD_600nm_ 0.1 in 96-well blocks with rhamnose maintained at 0% or 0.05%. Once the uninduced cultures reached OD_600nm_ 0.3, 120 μL from all samples was mixed with 3X RNAgem Blue Buffer (Zygem, Charlottesville VA), and chemically lysed by incubation at 75 °C for 10 minutes in a thermocycler.

### Generation of RNA-Seq data

Illumina cDNA libraries were generated using a modified version of the RNAtag-seq protocol.^59^ Briefly, 500ng-1 μg of total RNA was fragmented, depleted of genomic DNA, dephosphorylated, and ligated to DNA adapters carrying 5’-AN_8_-3’ barcodes of known sequence with a 5’ phosphate and a 3’ blocking group. Barcoded RNAs were pooled and depleted of rRNA using the RiboZero rRNA depletion kit (Epicentre). Pools of barcoded RNAs were converted to Illumina cDNA libraries in 2 main steps: (i) reverse transcription of the RNA using a primer designed to the constant region of the barcoded adaptor with addition of an adapter to the 3’ end of the cDNA by template switching using SMARTScribe (Clontech) as described^60^; (ii) PCR amplification using primers whose 5’ ends target the constant regions of the 3’ or 5’ adaptors and whose 3’ ends contain the full Illumina P5 or P7 sequences. cDNA libraries were sequenced on the Illumina [NextSeq 500] platform to generate paired end reads.

### Computational analysis of RNA-Seq data

Sequencing reads from each sample in a pool were demultiplexed based on their associated barcode sequence using custom scripts. Up to 1 mismatch in the barcode was allowed provided it did not make assignment of the read to a different barcode possible. Barcode sequences were removed from the first read as were terminal G’s from the second read that may have been added by SMARTScribe during template switching.

### Pathway analysis of RNA-Seq data

DESeq2^61^ was used to calculate the differential gene expression (log_2_ fold change and adjusted p-values) for each CRISPRi strain relative to the control-CRISPRi strain at matched rhamnose dose or time point.

Pathways enriched with significantly up or downregulated genes with an adjusted p-value cutoff of 0.05 were identified using STRINGdb^62^ and visualized using Morpheus^63^.

### PerSpecTM analysis

Target prediction using the CRISPRi^R^ P_rhaBAD_ strains was performed as previously described.^40^ Briefly, z-scored gene count tables were used to calculate Pearson correlation values between CRISPRi^R^ strain knockdown and the PA14 reference strain treated with compound for 90 minutes. Gene expression profiles from fosmidomycin treatment used a *dxr*-hypomorph as fosmidomycin does not have wild type PA14 activity. A target correlation score (*r̅_max_*) was calculated between compound (as the query) and CRISPRi^R^ strain (as the reference set). The correlation scores were visualized using Morpheus.

### Genomic DNA extraction and Illumina sequencing

Genomic DNA (gDNA) was isolated from bacterial sweeps from pooled conjugation selective LB agar plates or from aliquots of culture from uninduced passaging of CRISPRi pools using the Qiagen DNeasy Blood & Tissue Kit (#69506). gDNA was quantified using the Qubit dsDNA kit (#Q32854, Invitrogen). The sgRNA region was amplified from 100 ng of gDNA using NEBNext Ultra II Q5 master mix (#M0544L, New England Biolabs, Ipswich MA) and the following PCR conditions: 98 °C for 45 s; 17 cycles of 98 °C for 10 s, 64 °C for 30 s, 65 °C for 20 s; 65 °C for 5 min. The sgRNAs were amplified using forward primer 13 (0.5 μM final concentration) that appends a P5 flow cell adapter sequence and a Read1 Illumina sequencing primer binding site, and reverse primer 14 (0.5 μM final concentration) that appends a P7 flow cell adapter sequence and a Read2 Illumina sequencing primer binding site. Primers contain unique barcodes for sample pooling during Illumina sequencing. The ∼230 bp amplicons were purified using AMPure XP beads (Beckman-Coulter #A63880) with one-sided selection (1.2X). Amplicon size, purity, and concentration were measured using D1000 DNA ScreenTape (#5067-5582, Agilent, Santa Clara CA) on a 2200 TapeStation (Agilent, Santa Clara CA). Amplicons from each sample were multiplexed sequenced with Illumina MiSeq 300 (Paired-end 2x150 cycles, 1x8 i5 cycles, and 1x8 i7 cycles).

### Quantification of sgRNA

To measure the abundance of sgRNAs from the pooled conjugations or from the CRISPRi pools, sgRNA sequencing read pairs were processed using Trimmomatic^64^ and aligned using subread-align^65^.

Trimmomatic clipped the bases before base 30 on the 5’ end and after base 80 on the 3’ end, generating ∼ 50bp reads. Processed 50bp reads were aligned to the full-length reference sgRNA sequence (∼100bp) using subread-align with default parameters, minFragLength = 30, maxFragLength = 100. For the pooled conjugations, the percentage of mapped sgRNA reads was calculated for each sgRNA relative to the total number of mapped sgRNA reads such that each sgRNA should have ∼ 5% of the total mapped sgRNA reads (18 strains for 100% of reads, 1 strain ∼ 5%). For the uninduced passaging of CRISPRi pools, the percentage of mapped sgRNA reads was calculated but reported as the log2-fold change of each guide relative to time zero.

## ACKNOWLEDGEMEMTS

We would like to thank Sophie Chen for assistance with code in R. RNA-Seq libraries were constructed and sequenced at the Broad Institute of MIT and Harvard by the Microbial ‘Omics Core and Genomics Platform, respectively. The Microbial ‘Omics Core also provided guidance on experimental design and conducted preliminary analysis for all RNA-Seq data. This research was supported by a U19 NIH grant (U19AI142780). All figures were created with BioRender.com.

## AUTHOR CONTRIBUTIONS

Conceptualization, J.R.S., T.W., D.T.H.; investigation J.R.S., K.F., R.B.; formal analysis J.R.S., K.F.; writing— original draft J.R.S.; writing—review & editing J.R.S.,K.F., R.B., K.P.R., T.W., D.T.H.; resources D.T.H.; supervision T.W., D.T.H.; funding acquisition D.T.H.

## DECLARATION OF INTEREST

The authors declare no competing interests.

## Notes

### Competing Interest Statement

The authors have declared no competing interest.

