## Supplemental figures for "An Inducible CRISPRi system for phenotypic analysis of essential genes in *Pseudomonas aeruginosa*"

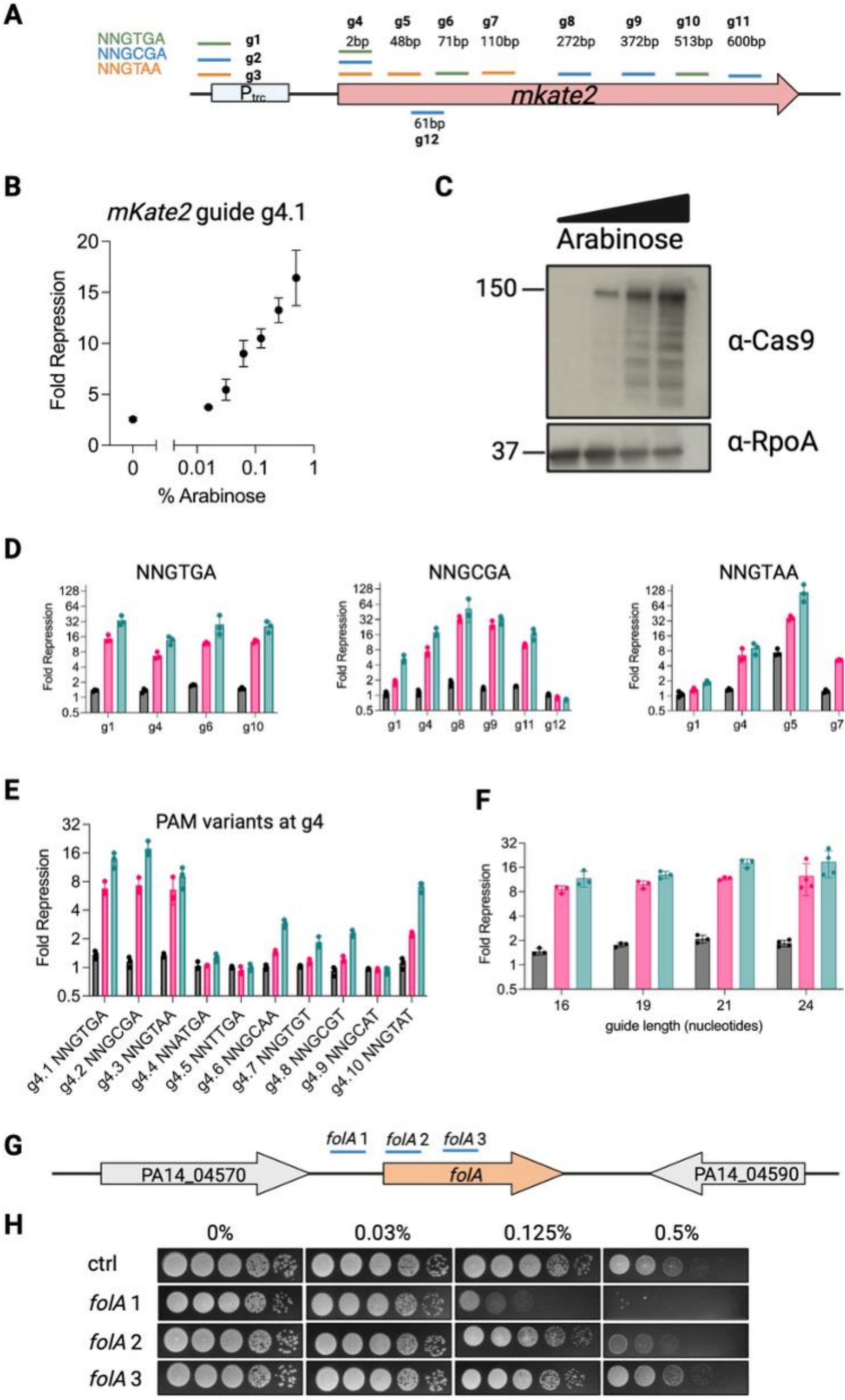
A
NNGTGA
NNGCGA
NNGTAA
g1
g2
g3
g4
2bp
g5
48bp
g6
71bp
g7
110bp
g8
272bp
g9
372bp
g10
513bp
g11
600bp
Ptrc
mkate2
61bp
g12
B
mKate2 guide g4.1
Fold Repression
% Arabinose
C
Arabinose
150
α-Cas9
37
α-RpoA
D
NNGTGA
NNGCGA
NNGTAA
Fold Repression
0%
0.125%
0.5%
E
PAM variants at g4
Fold Repression
g4.1 NNGTGA
g4.2 NNGCGA
g4.3 NNGTAA
g4.4 NNATGA
g4.5 NNTTGA
g4.6 NNGCAA
g4.7 NNGTGT
g4.8 NNGCGT
g4.9 NNGCAT
g4.10 NNGTAT
F
Fold Repression
guide length (nucleotides)
0%
0.125%
0.5%
G
folA 1
folA 2
folA 3
PA14_04570
folA
PA14_04590
H
0%
0.03%
0.125%
0.5%
ctrl
folA 1
folA 2
folA 3

Figure S1. Characterization of the arabinose-based $CRISPRi^A$ system. (A) Schematic of the open reading frame of the fluorescent reporter mKate2 with PAMs and sgRNA locations highlighted. Base pairs indicate distance from *mKate2* start codon. (B) mKate2 fluorescence repression (normalized to control sgRNA) at varying arabinose doses using sgRNA g4.1 that targeted the start codon of *mKate2*. Fluorescence reduction showed direct correlation with arabinose concentration and thus $dCas9_{Spa}$ expression levels. (C) Western blot probing for $dCas9_{Spy}$ expression with a dCas9-specific antibody after induction indicated protein degradation at higher arabinose doses. (D) Effect of varying guide location with the three reported $dCas9_{Spa}$-specific PAM sequences on mKate2 fluorescence repression. The guides bound to the promoter or open reading frame of *mKate2* as indicated in panel A. NNGTGA and NNGCGA were equally effective irrespective of guide location while NNGTAA was weaker and displayed location bias. (E) Impact of PAM sequence engineered in the start codon-targeting sgRNA g4 on $dCas9_{Spa}$ activity as measured by mKate2 fluorescence repression. NNGTGA and NNGCGA led to the highest mKate2 depletion (16-fold fluorescence repression) and were closely followed by NNGTAA and the newly designed NNGTAT (~8-fold repression). (F) Impact of length of the targeting region of the guide. Targeting region length between 16 and 24 nucleotides (nt) showed similar extent of fluorescence repression. (G) Schematic of guide design to target *folA* using $CRISPRi^A$. sgRNAs were designed to target the promoter region (*folA1*) or the coding sequence (*folA2*, *folA3*). (H) Growth defects from *folA* knockdown. Spot dilutions of $CRISPRi^A$ strains with control or *folA* sgRNAs grown on LB agar with varying arabinose concentrations. Growth defects were more apparent when the promoter region was targeted.

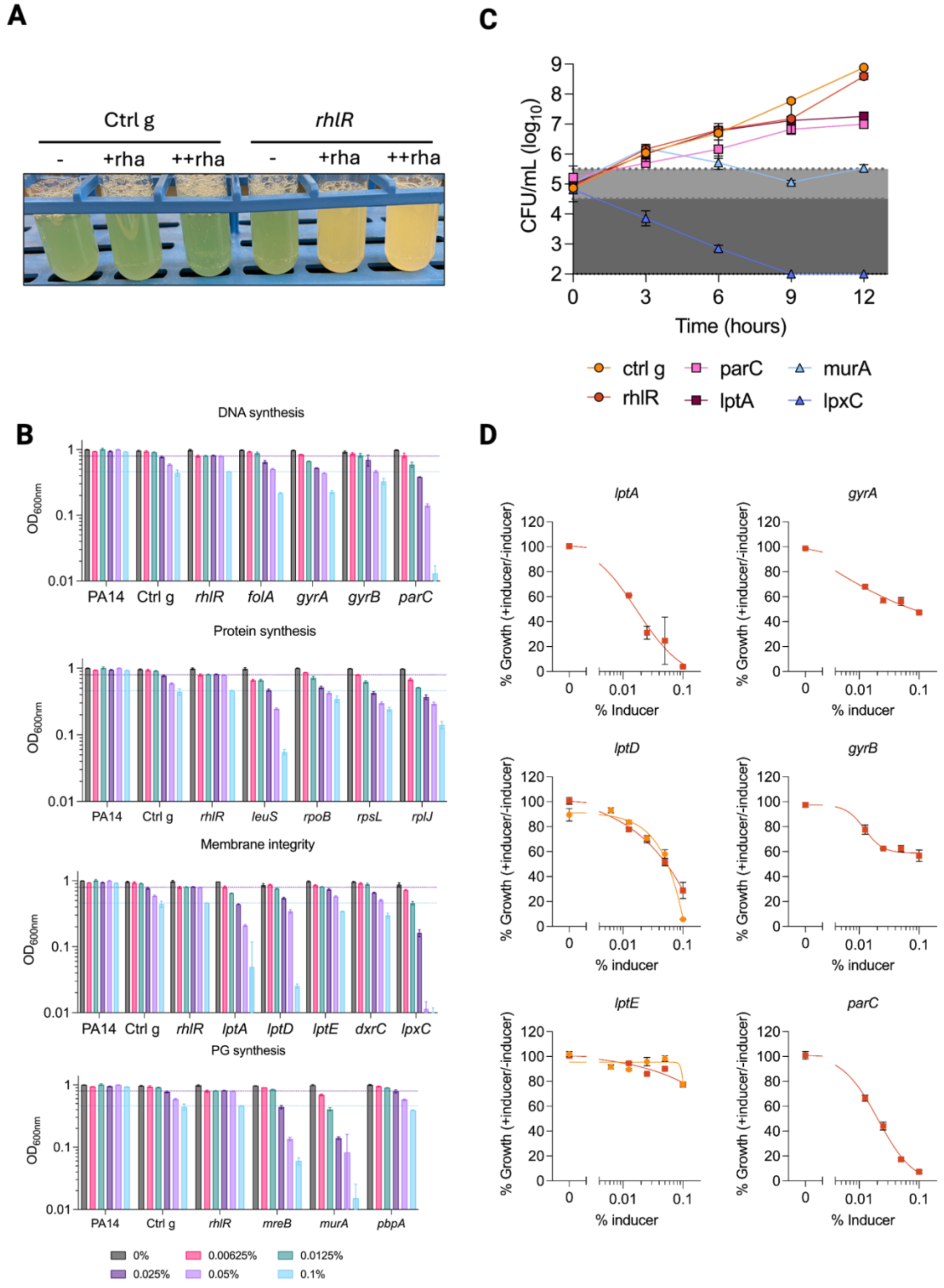


Figure S2. Phenotypic effects of CRISPRi$^{R}$ knockdown. (A) CRISPRi$^{R}$ was effective against endogenous non-essential gene, *rhlR*, but had minimal impact on growth. Knockdown with 0.0125% rhamnose (+) or 0.05% rhamnose (++) exposure overnight caused significant reduction of pigmented pyocyanin, whose production is regulated by RhlR, relative to the *control*-CRISPRi strain. (B) $CRISPRi^R$ targeting of endogenous essential genes led to rhamnose dose-dependent growth inhibition. Individual CRISPRi strains were grown in LB broth in the presence of 0%, 0.00625%, 0.0125%, 0.025%, 0.05% or 0.01% rhamnose and optical density at 600nm ($OD_{600nm}$) was measured after 24 hours. Dashed lines indicate $OD_{600nm}$ after knockdown of the non-essential gene *rhlR*. (C) Kill kinetics of $CRISPRi^R$ strains grown in LB with 0.05% rhamnose. Effect of knockdown on growth is shown with background colour (white – slowed growth, light grey – bacteriostatic, dark grey – bactericidal). (D) Dose-response curves of the growth of $CRISPRi^R$ strains targeting PA14 genes involved in LPS transport or related to DNA synthesis at varying concentrations of the inducer, rhamnose. $CRISPRi^R$ strains were grown in LB broth with varying concentrations of rhamnose for 24 hours and $OD_{600nm}$ was read to measure growth. It was normalized to uninduced conditions and plotted as %Growth. The degree of growth defect is gene-specific and pathway-independent.

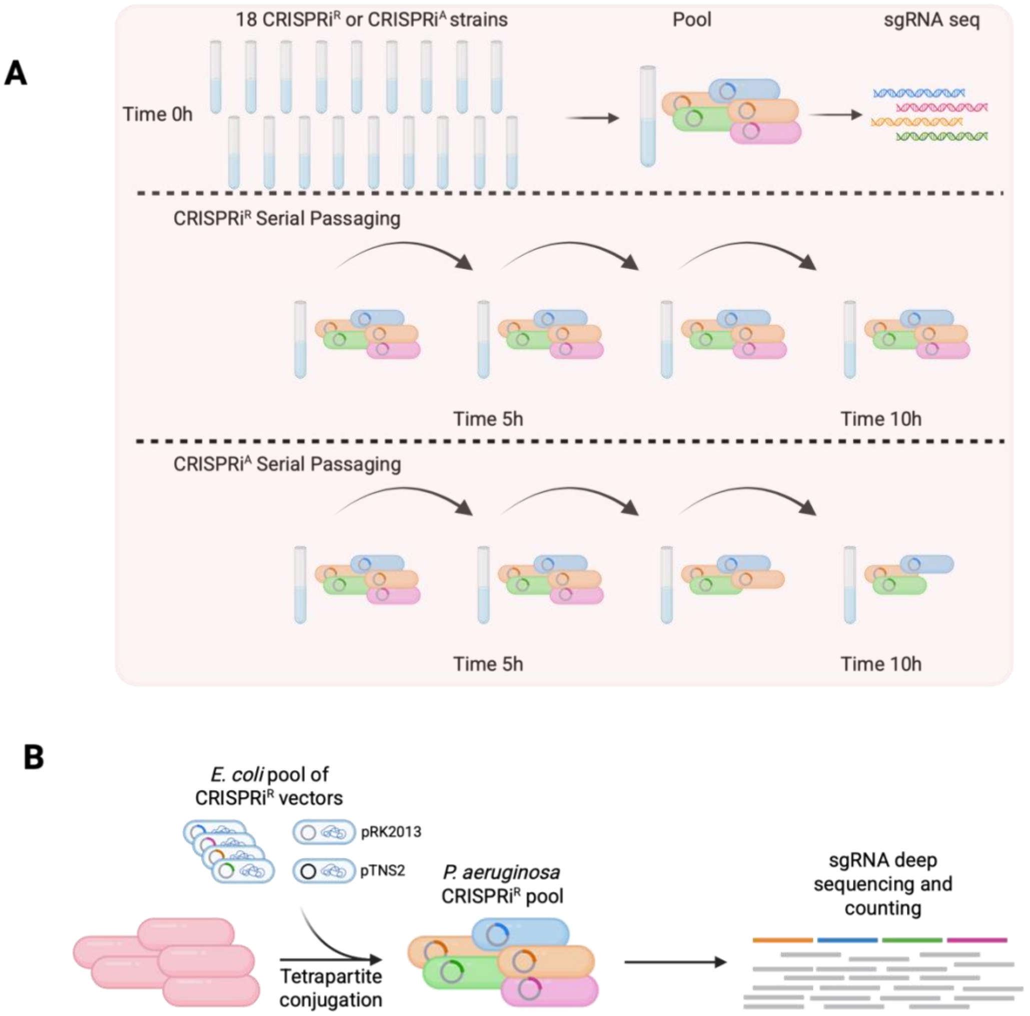


Figure S3. CRISPRi$^R$ facilitates pooled CRISPRi library construction in *P. aeruginosa*. (A) Schematic showing the propagation of CRISPRi pools without inducer. All CRISPRi strains were grown arrayed and then pooled at equal cell densities. Pooled strains were grown from $OD_{600}$ 0.025 until mid-log and passaged back to $OD_{600}$ 0.2 for 5 cycles. Aliquots were collected after 0, 5, and 10 hours of passaging, gDNA extracted and processed for sgRNA-targeted amplicon sequencing. (B) Schematic showing the pooled CRISPRi conjugation strategy. The tetrapartite mating is set up between recipient PA14::*dcas9*, *E. coli* helper pRK2013, *E. coli* helper pTNS2, and a pool of *E. coli* donor CRISPRi$^R$ strains (17 different *E. coli* CRISPRi$^R$ sgRNA strains pooled at equal cell density). After selection of true PA14 CRISPRi$^R$ conjugants with irgasan (5 µg/mL) and gentamicin (15 µg/mL), colonies were pooled together, gDNA extracted and processed for sgRNA-targeted amplicon sequencing.

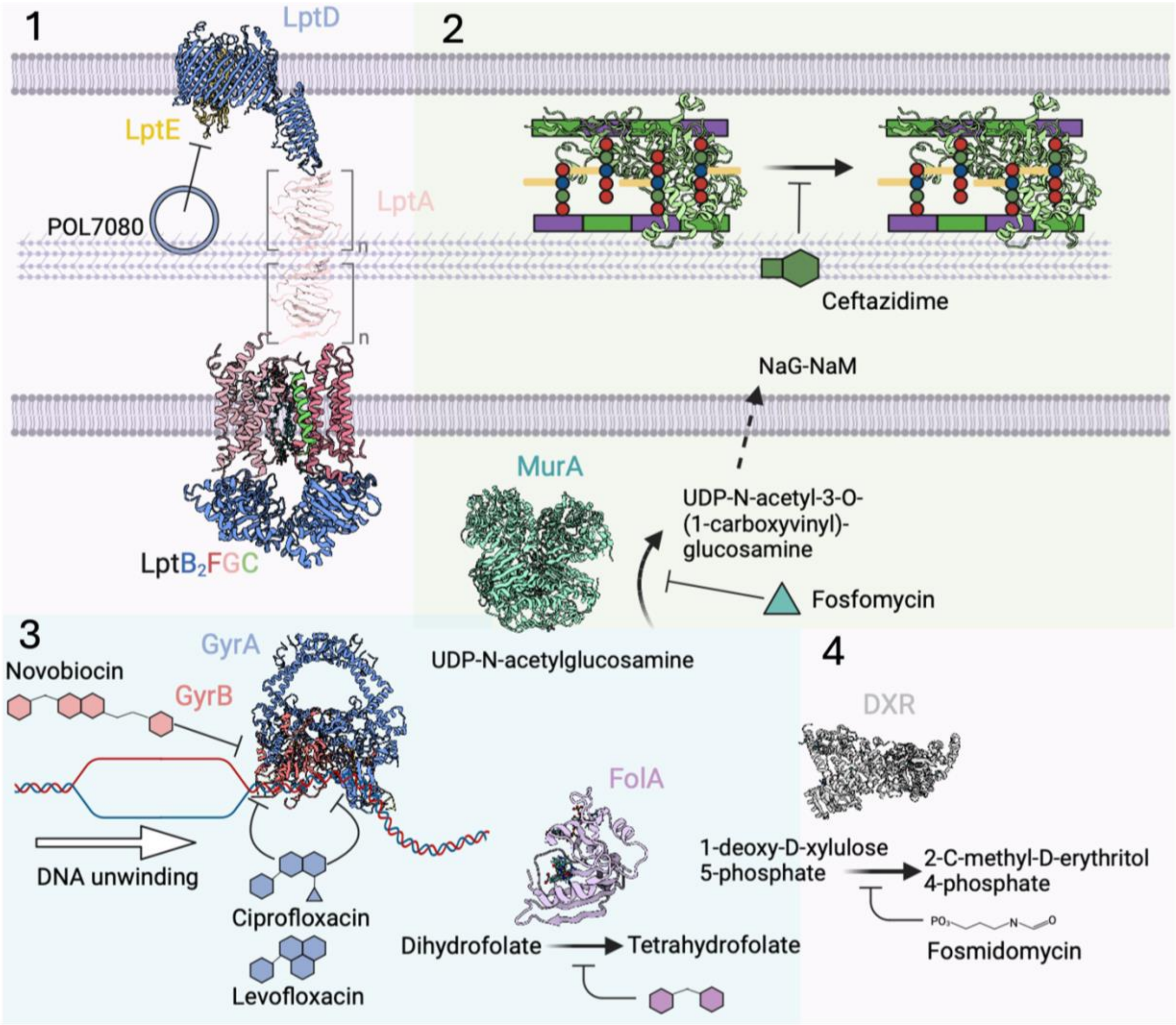


Figure S4. Cartoon of chemical inhibitors and their targets in (1) LPS trafficking, (2) peptidoglycan synthesis, (3) DNA synthesis-related, and (4) dxr pathways in *P. aeruginosa*.

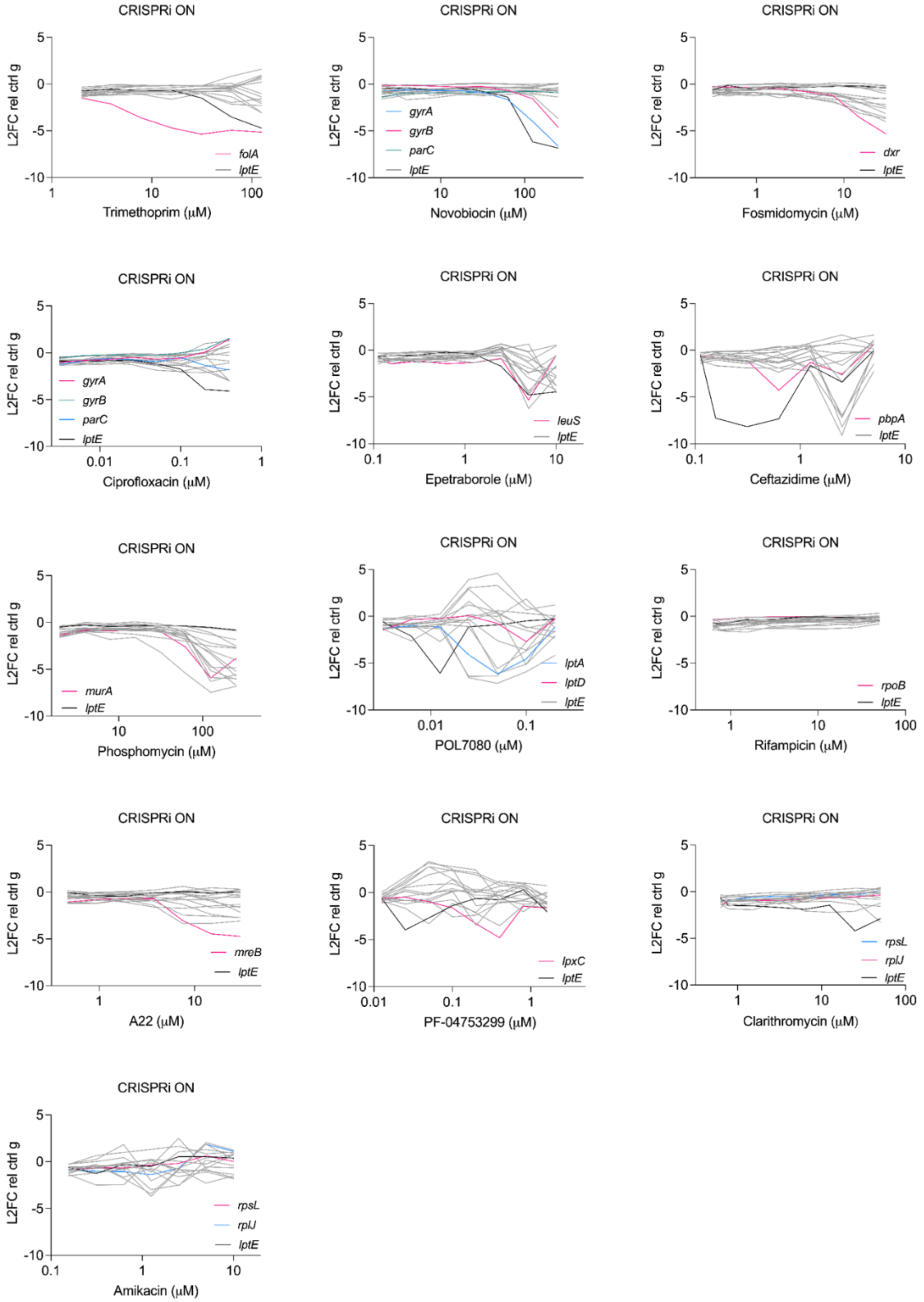
CRISPRi ON
L2FC rel ctrl g
folA
lptE
Trimethoprim (μM)
gyrA
gyrB
parC
lptE
Novobiocin (μM)
dxr
lptE
Fosmidomycin (μM)
gyrA
gyrB
parC
lptE
Ciprofloxacin (μM)
leuS
lptE
Epetraborole (μM)
pbpA
lptE
Ceftazidime (μM)
murA
lptE
Phosphomycin (μM)
lptA
lptD
lptE
POL7080 (μM)
rpoB
lptE
Rifampicin (μM)
mreB
lptE
A22 (μM)
lpxC
lptE
PF-04753299 (μM)
rpsL
rplJ
lptE
Clarithromycin (μM)
rpsL
rplJ
lptE
Amikacin (μM)

Figure S5. Dose-response curves of the $log_2$ fold change (L2FC) in growth of CRISPRi$^R$ strains relative to *control*-CRISPRi$^R$ strain. CRISPRi$^R$ strains were grown in sub-lethal rhamnose doses. *folA*-, *gyrA*-, *gyrB*-, *lptE*-, *rpoB*-, and *pbpA*-CRISPRi strains were grown in 0.05% rhamnose. *parC*-, *lptA*-, *lptD*-, *dxr*-, *mreB*-, *leuS*-, *rpsL*, and *rplJ*-CRISPRi strains were grown in 0.025% rhamnose. *lpxC*- and *murA*-CRISPRi strains were grown in 0.0125% rhamnose. Growth was measured with optical density at 600 nm after 24 hours in the presence of chemical inhibitor. Fold change in growth relative to *control*-CRISPRi strain grown with identical rhamnose doses is plotted. The putative genetic target of the chemical inhibitor is highlighted in pink. *gyrB* and *parC* are highlighted as non-binding targets and alternative targets, respectively for ciprofloxacin. *lptA* and *lptE* are highlighted as on pathway targets for POL7080. *rplJ* and *rpsL* are highlighted as on pathway targets for amikacin and clarithromycin. *lptE*-CRISPRi strain is broadly sensitized to many chemical inhibitors and is highlighted for all chemical inhibitors.

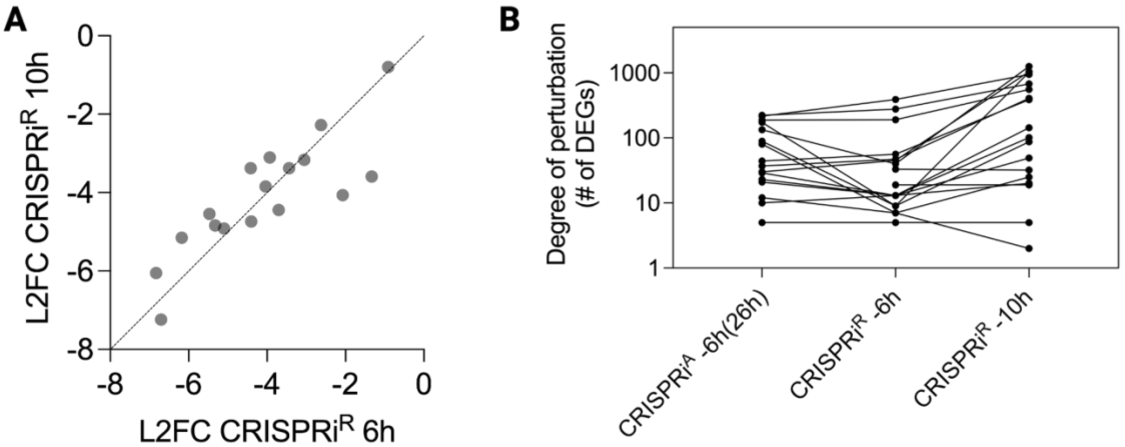


Figure S6. Tighter control and extended CRISPRi$^R$ activation results in greater transcriptomic perturbation. (A) CRISPRi$^R$ knockdown analyzed by RNA-seq and quantified using $log_2$ fold change (L2FC) of target gene relative to *control*-CRISPRi strain after 6 and 10 hours of growth in 0.05% rhamnose. Each point is an individual CRISPRi$^R$ strain. Measuring L2FC at 10 hours required a modified experimental set up: strains were grown in LB 0.05% rhamnose from $OD_{600nm}$ 0.05 to 0.5 (first 6 hours) and back diluted to $OD_{600nm}$ 0.1 and allowed to grow until $OD_{600nm}$ 0.3 (an additional 4 hours). Back dilution of the CRISPRi$^R$ strains prevented entry into stationary phase for accurate transcriptional profiling at the 10-hour time point. (B) Differentially expressed genes (DEGs) from target gene knockdown using CRISPRi$^A$ after 6 hours of induction with 0.5% arabinose (~effectively 26 hours of induction due to the leaky system) or CRISPRi$^R$ after 6 and 10 hours of induction with 0.05% rhamnose. The extended induction of 10 hours with CRISPRi$^R$ captured a more comprehensive transcriptional perturbation downstream of target depletion for most genes as reflected in the increase in DEGs relative to CRISPRi$^A$ or the shorter 6-hour induction with CRISPRi$^R$.

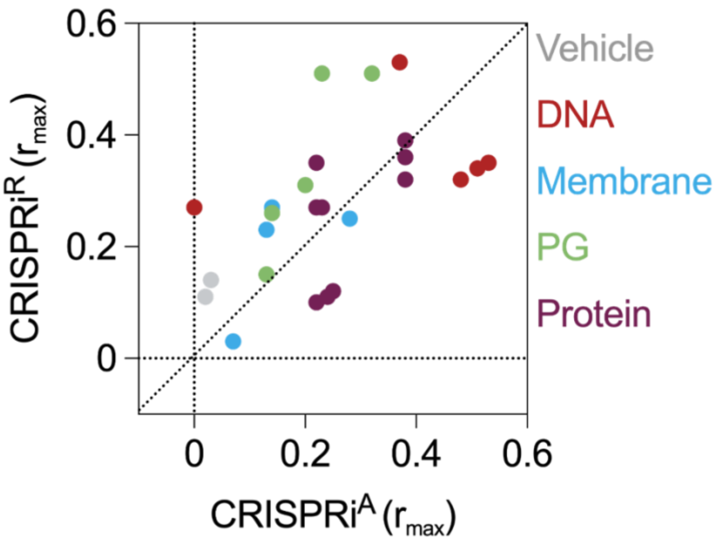


Figure S7. Pearson correlations from PerSpecTM analysis of chemical inhibitors queried against CRISPRi strains are higher from CRISPRi$^{R}$ target knockdown. Maximum Pearson correlations ($r_{max}$) for each chemical inhibitor and its known target knockdown using each CRISPRi system are shown and color coordinated by mechanism of action (Vehicle: water, DMSO; DNA: ciprofloxacin, levofloxacin, cisplatin, novobiocin, trimethoprim; Membrane: fosmidomycin, PF3299, POL7080, colistin; PG: A22, phosphomycin, carbenicillin, imipenem, ceftazidime; Protein: rifampicin, amikacin, gentamicin, tobramycin, tetracycline, doxycycline, minocycline, clarithromycin, erythromycin). CRISPRi$^{R}$ $r_{max}$ values are from the heatmap in Figure 5A; CRISPRi$^{A}$ $r_{max}$ values are from Romano, K. P. *et al.* Perturbation-specific transcriptional mapping for unbiased target elucidation of antibiotics. *Proc Natl Acad Sci U S A* **121**, (2024).
